# A bi-directional binding site linking the α_2_δ-1 subunit to the intrinsic speed control process in VSD I of voltage-gated calcium channels

**DOI:** 10.1101/2025.10.07.680875

**Authors:** Martin C. Heiss, Monica L. Fernández-Quintero, Nicole Kranbitter, Hajar El Aouad, Marta Campiglio, Bernhard E. Flucher

**Affiliations:** Institute of Physiology, Department of Physiology and Medical Biophysics, Medical University Innsbruck, 6020 Innsbruck, Austria; Institute of General, Inorganic and Theoretical Chemistry, University of Innsbruck, 6020 Innsbruck, Austria

**Author notes:** Correspondence to: Bernhard E. Flucher, PhD Institute of Physiology, Department of Physiology and Medical Physics Medical University of Innsbruck 6020 Innsbruck, Austria.

## Abstract

Voltage-gated calcium channels communicate electrical signals in membranes of excitable cells into cellular responses like secretion of hormones and neurotransmitters, or the contraction of heart and skeletal muscle cells. Their activation properties are tuned to match their specific functions. Consequently, the different members of the calcium channel family activate over a wide range of voltages and with greatly differing speeds. The skeletal muscle Ca_V_1.1 and the cardiac/neuronal Ca_V_1.2 represent two structurally closely related channels with particularly slow and fast activation kinetics, respectively. Both channel paralogs associate with the auxiliary calcium channel subunit α_2_δ-1, which is a known regulator of activation properties. By expressing Ca_V_1.1 and Ca_V_1.2 with and without α_2_δ-1 in a new double-knockout muscle cell line, we demonstrate that α_2_δ-1 regulates activation kinetics of the two channels in opposite directions. Molecular dynamics simulation revealed a string of charged amino acids connecting α_2_δ-1 to the intrinsic speed-control mechanism of voltage-sensing domain I (VSD I) in Ca_V_1.1. Charge-neutralizing mutations of any of these charged amino acids abolished the α_2_δ-1 modulation and accelerated current kinetics. Together, these results reveal the molecular mechanism by which the α_2_δ-1 subunit regulates the intrinsic speed-control mechanism in the VSD I of Ca_V_1.1 calcium channels.

## Introduction

High voltage-gated calcium channels are multi-subunit membrane proteins critically involved in the regulation of vital cell functions such as heart and skeletal muscle contraction, secretion of hormones and neurotransmitters, and gene regulation. To accomplish these diverse tasks, the members of the Ca_V_ channel family exhibit a broad range of gating properties, thus conducting calcium currents that activate at greatly different voltages and with distinct kinetics.

In this regard, assembly of macromolecular complexes of the pore-forming α_1_ subunit with the auxiliary α_2_δ, β and γ subunits is important to “normalize” the current properties; i.e. to provide a channel with the voltage dependence and kinetics characteristic for its currents in the native cells. Specifically, the α_2_δ-1 subunit is known to regulate the activation kinetics of L-type calcium channels. In previous siRNA knockdown experiments, we demonstrated that in muscle cells the auxiliary α_2_δ-1 subunit supports the intrinsically slow and fast activation kinetics of the skeletal muscle and cardiac calcium channels, respectively; making Ca_V_1.1 currents slow and Ca_V_1.2 currents fast ^1–3^.

The calcium channel α_2_δ-1 subunit is a highly glycosylated extracellular protein, tethered to the membrane by a GPI-anchor and contacting the α_1_ subunit at multiple extracellular sites. The cryo-EM structure of the skeletal muscle calcium channel complex identified the first extracellular loop of VSD I (designated L1-2_I_) as the interaction site between Ca_V_1.1 and the functionally important metal ion-dependent adhesion site (MIDAS) of α_2_δ-1 ^4,5^. In Ca_V_1.2 negatively charged residues in the corresponding L1-2_I_ loop of the first voltage-sensing domain (VSD I) were shown to be of critical importance for the association of α_2_δ-1 and current modulation by α_2_δ-1 ^6^.

In Ca_V_1.1 and Ca_V_1.2, VSD I determines the channeĺs specific activation kinetics. Molecular dynamic (MD) simulations of Ca_V_1.1 indicated that, when an electric potential is applied, VSD I undergoes state-transitions between the activated and resting states considerably slower than the three other VSDs ^7^. Voltage-clamp fluorometry recordings of Ca_V_1.2 demonstrated that co-expression of α_2_δ-1 specifically increased the kinetics of VSD I ^8^. Together, these observations suggest that VSD I is rate limiting for the slow activation kinetics of Ca_V_1.1 and the comparably fast kinetics of Ca_V_1.2. Notably, these properties can be exchanged between the two channel isoforms by swapping sequences containing the IS3 helix and IS3S4 loop ^9,10^. Furthermore, recent mutagenesis studies in Ca_V_1.1 identified the role of interactions between the positive S4 gating charges and negative countercharges in the IS2 helix in determining the Ca_V_1.1 activation kinetics. Specifically, charge-neutralizing mutations of the outermost gating charge K0 and of a countercharge E90 in the extracellular part of IS2 resulted in the acceleration of activation kinetics ^11,12^. Therefore, the intrinsic molecular machinery controlling the specific kinetic properties of Ca_V_1 channels depends on the interaction of highly conserved charged residues in IS2 and IS4, as well as on isoform-specific structural features of IS3 and the IS3-S4 loop.

Altogether, these findings suggest that both, intrinsic structural features of VSD I and the action of the auxiliary α_2_δ-1 subunit cooperate in controlling the speed of L-type calcium channels. Yet, how the two mechanisms are connected with each other remains incompletely understood. Here, we used site-directed mutagenesis of a putative interaction site between α_2_δ-1 and VSD I of Ca_V_1.1 to reveal a direct molecular link between the intrinsic and extrinsic regulators of activation kinetics. Our data indicate that, via a cluster of negatively charged amino acids in the L1-2_I_ loop of VSD I, the α_2_δ-1 orchestrates a network of ionic interactions, which regulate the speed of VSD I state transitions that in turn control Ca_V_1.1 activation kinetics.

## Results

### Knockout of α_2_δ-1 in muscle cells changes current kinetics of Ca_V_1.1 and Ca_V_1.2 in opposite directions

Previously, we demonstrated that siRNA knockdown of α_2_δ-1 in dysgenic myotubes reconstituted with Ca_V_1.1 accelerates current kinetics, whereas in myotubes expressing Ca_V_1.2 it decelerates current kinetics ^1–3^. This suggested that the function of the α_2_δ-1 subunit is to stabilize the intrinsic current properties of the respective pore-forming α_1_ subunit. Here we re-visited this issue by generating a new muscle expression system lacking both Ca_V_1.1 and α_2_δ-1. Starting from the dysgenic (Ca_V_1.1-null) cell line GLT, we used the CRISPR-Cas9 technology to additionally knock out expression of the α_2_δ-1 subunit (**Fig. 1A**). Consistent with the genotyping results, cells of the selected clone C9 showed a strong reduction in α_2_δ-1 mRNA levels and lacked detectable α_2_δ-1 protein expression (**Fig. 1B-C**). When we reconstituted this double Ca_V_1.1/α_2_δ-1 knockout cell line (designated GLT-C9) with Ca_V_1.1 and α_2_δ-1, both channel subunits co-clustered with the type 1 ryanodine receptor (RyR1) (**Fig. 1D**). In GLT myotubes and the cell lines derived thereof these clusters represent calcium release units primarily formed between the SR and the plasma membrane (peripheral junctions) and to a lesser degree with transverse (T-) tubules (triads) ^13,14^. For simplicity reasons and in line with the terminology used in mature muscle, henceforth we will subsume these functionally equivalent junctions under the term “triad” and the correct insertion of channels into these junctions as “triad targeting”. Thus, recombinant Ca_V_1.1 and α_2_δ-1 co-aggregate together in triad junctions of GLT-C9 myotubes and restore calcium currents (see below). When only one or the other channel subunit was expressed in GLT-C9 myotubes, Ca_V_1.1 still assumed a clustered distribution, but α_2_δ-1 remained diffusely expressed in the plasma membrane (**Fig. 1F-G**); like the native α_2_δ-1 in the mother GLT cell line ^1^. These data confirm our earlier findings showing that functional membrane expression and targeting of calcium channel α_1_ subunits into SR-plasma membrane junctions in skeletal muscle cells is independent of the presence of α_2_δ-1 ^1–3^. And conversely that triad targeting of α_2_δ-1, but not its membrane expression, depends on the presence of an α_1_ subunit ^15^. Furthermore, we reconstituted the GLT-C9 cells with the Ca_V_1.2 isoform plus α_2_δ-1 (**Fig. 1E**). Also this combination of channel subunits co-localized in the typical clusters, indicating that also the cardiac/neuronal Ca_V_1.2 subunit is targeted into the triad junctions of the GLT-C9 myotubes ^16^. Again, triad targeting of Ca_V_1.2 did not require the presence of the α_2_δ-1 subunit.

**Fig. 1.**
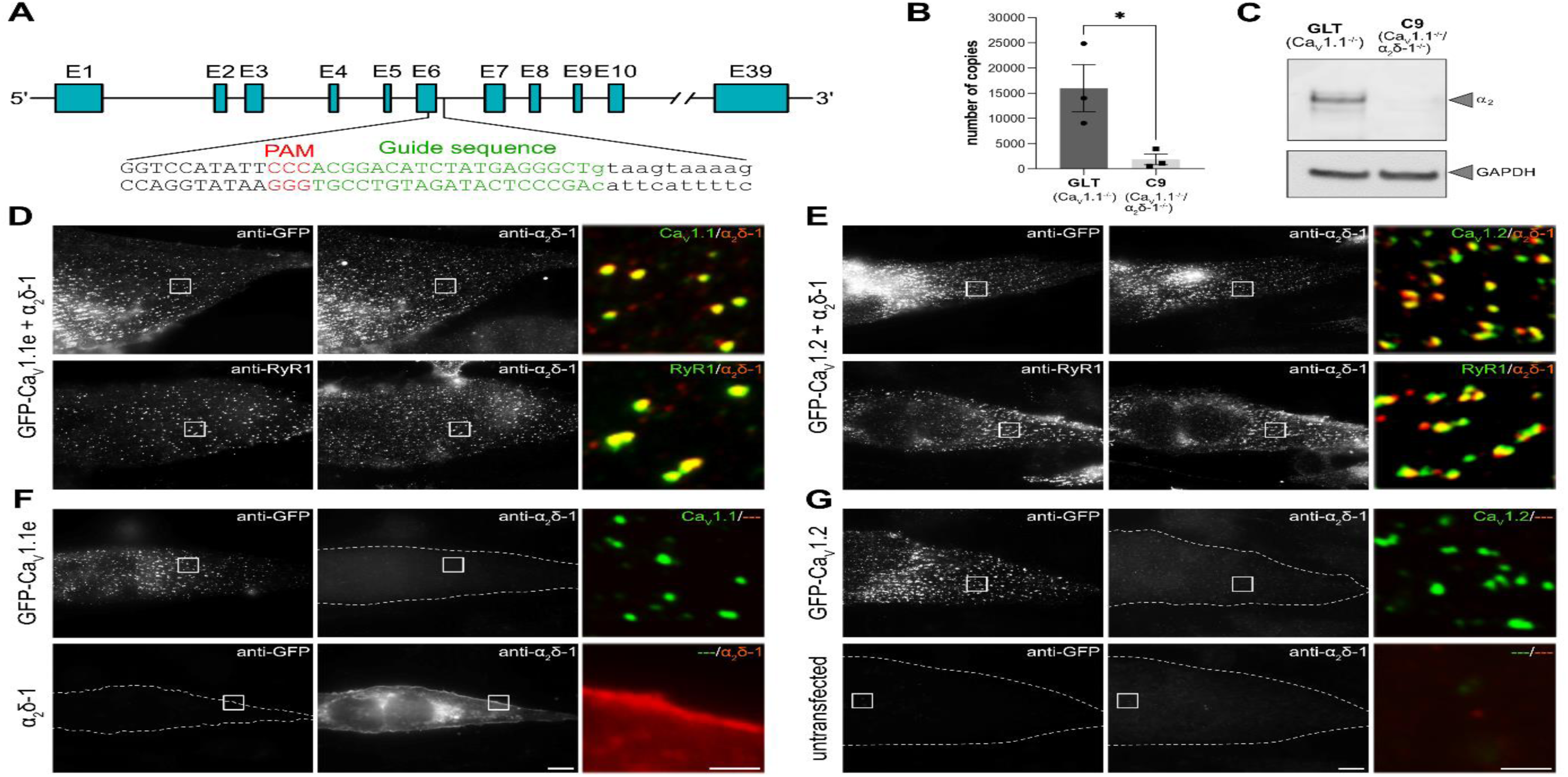
Characterization of GLT-C9 (Ca_V_1.1/α_2_δ-1-null) myotubes and triad targeting of Ca_V_1.1 and Ca_V_1.2 with and without α_2_δ-1. **A**: Diagram of the α_2_δ-1 locus with exons depicted. Exon 6 of α_2_δ-1 was targeted in GLT cells using a vector encoding the specified guide sequence RNA and Cas9. **B**: Quantification of α_2_δ-1 mRNA levels in GLT cells and the selected C9 α_2_δ-1-KO clone by TaqMan quantitative PCR. Data represent the mean ± SEM of three independent cDNA preparations. Differences between groups are statistically significant (P = 0.042). **C**: Western blot analysis (representative of six experiments) using the anti-α_2_δ-1 antibody 20A detected bands around 150 kDa in GLT cells, which were absent in C9 cells. The presence of multiple bands likely reflects different glycosylation states of α_2_δ-1. **D-G:** Immunofluorescence labeling of Ca_V_1.1 and Ca_V_1.2 expressed in GLT-C9 myotubes with and without α_2_δ-1. **D:** Co-expression of Ca_V_1.1 and α_2_δ-1 in GLT-C9 cells resulted in the co-assembly of both calcium channel subunits in clusters co-localized with RyR1; indicative of correct triad targeting of Ca_V_1.1 and α_2_δ-1. **E:** Co-expression of Ca_V_1.2 with α_2_δ-1 resulted in similar co-clustering with RyR1 in triads. **F,G:** Ca_V_1.1 or Ca_V_1.2 expressed without α_2_δ-1 in GLT-C9 myotubes still clustered in triads. α_2_δ-1 expressed without a Ca_V_1 α_1_-subunit (**F, lower**) became diffusely distributed in the cell membrane. Untransfected GLT-C9 cells (**G, lower**) expressed neither a Ca_V_1 subunit nor α_2_δ-1. Scale bar, 10 µm; merged color image, 10x magnification of areas indicated by box, scale bar, 2µm.

Importantly, patch-clamp analysis of calcium currents in transfected GLT-C9 myotubes showed that co-expression of α_2_δ-1 with either Ca_V_1.1 or Ca_V_1.2 modulated the activation kinetics of the calcium currents in specific ways (**Fig. 2**). In the absence of α_2_δ-1, Ca_V_1.1 calcium currents in GLT-C9 cells activated with a time constant (τ) of 6 ms. On co-expression of Ca_V_1.1 with α_2_δ-1 current activation was slowed down by more than 3-fold, with a τ of 20 ms (**Fig. 2 A-C**). On the contrary, Ca_V_1.2 calcium currents in GLT-C9 cells transfected without the α_2_δ-1 subunit activated with a τ of 4.5 ms, but experienced a 3-fold acceleration, with a τ of 1.5 ms, when Ca_V_1.2 was co-expressed with α_2_δ-1 (**Fig. 2 D-F**). For Ca_V_1.1 the current density and voltage-dependence of activation was not significantly altered on co-expression of α_2_δ-1. However, for Ca_V_1.2, co-expression of α_2_δ-1 also shifted the voltage-dependence of activation by about 22 mV to more hyperpolarized test potentials (**Fig. 2 H,I**; **Table 1**). These results demonstrate that without α_2_δ-1, the activation kinetics of both, Ca_V_1.1 and Ca_V_1.2, are in the intermediate range and hardly differ from one another. Only in complex with the α_2_δ-1 subunit the two L-type calcium channel isoforms express their characteristic slow and fast activation kinetics (**Fig. 2G**). Thus, in muscle cells, the α_2_δ-1 subunit primarily functions as regulator of current kinetics and its role therein is not to make channel activation generally faster or slower, but to facilitate the intrinsic activation properties of the respective channel isoform; making Ca_V_1.1 slow and Ca_V_1.2 fast.

**Fig. 2.**
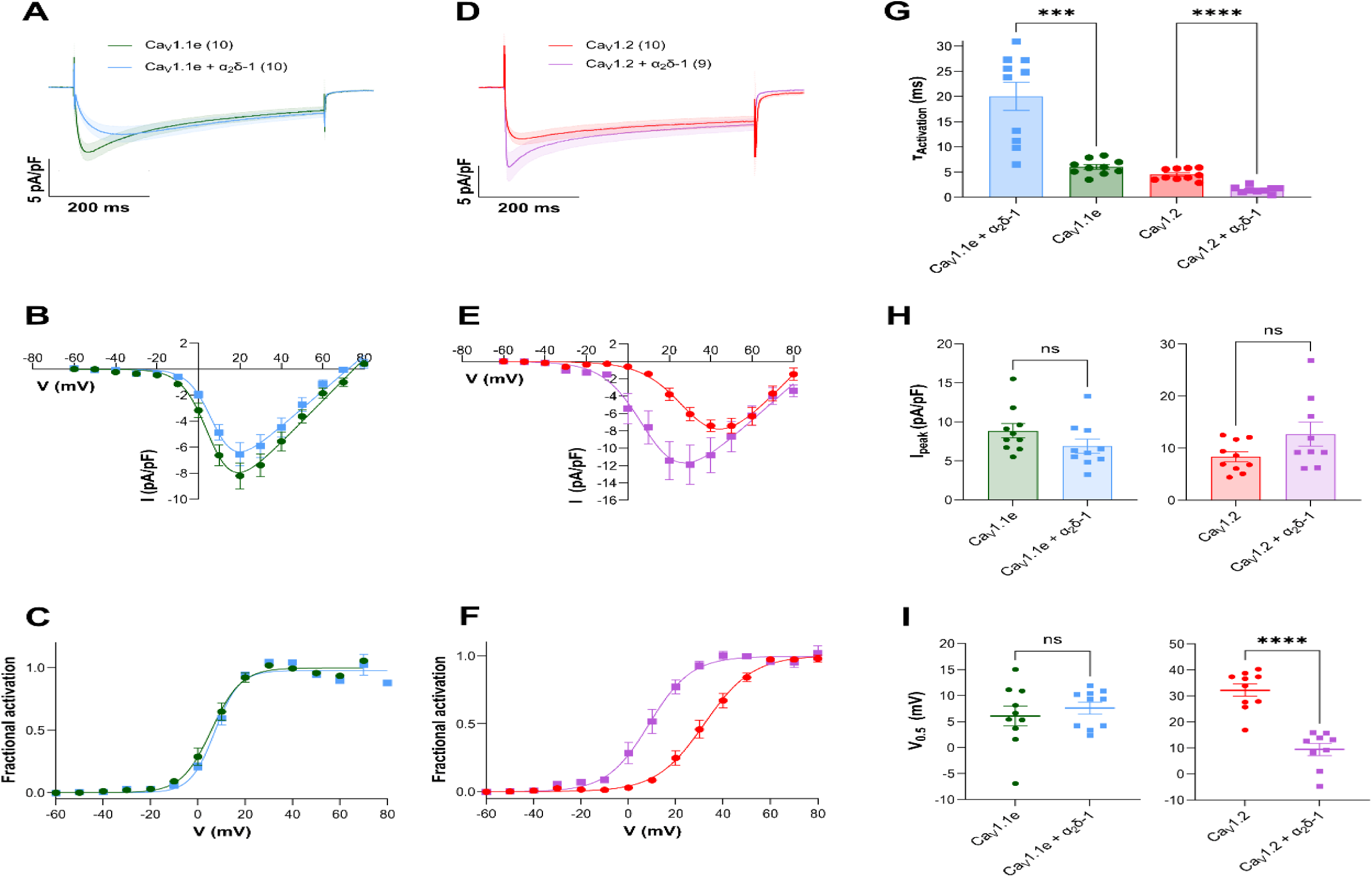
α_2_δ-1 functions as bi-directional regulator of current activation kinetics in GLT-C9 myotubes reconstituted with Ca_V_1.1 or Ca_V_1.2. GLT-C9 myotubes were reconstituted with either Ca_V_1.1 (**A-C**) or Ca_V_1.2 (**D-F**), with and without α_2_δ-1. **A,D**: Whole-cell calcium currents show that co-expression of α_2_δ-1 causes a substantial deceleration of activation kinetics in Ca_V_1.1 and a substantial acceleration in Ca_V_1.2. Voltage-dependence of activation was not affected by co-expression of α_2_δ-1 and Ca_V_1.1 (**C**) but was left-shifted to less positive potentials when α_2_δ-1 was co-expressed with Ca_V_1.2 (**F**). **G**: The time constants of current activation were very similar for Ca_V_1.1 and Ca_V_1.2 in the absence of α_2_δ-1, but assumed their characteristic slow and fast values on co-expression of α_2_δ-1. **H,I**: Quantitative comparison of peak current densities and voltage-dependence of activation reveal statistically significant effects of α_2_δ-1 co-expression only in the V½ of Ca_V_1.2 current activation. The data points are represented as mean ± SEM. Statistical tests: panel G, Welch’s t-test (Ca_V_1.1) and t-test (Ca_V_1.2); panel H, Mann-Whitney test (Ca_V_1.1) and Welch’s t-test (Ca_V_1.2); panel I, t-test (Ca_V_1.1) and Mann-Whitney test (Ca_V_1.2). ∗ ≜ *p* < 0.05, ∗∗ ≜ *p* < 0.01, ∗∗∗ ≜ *p* < 0.001, ∗∗∗∗ ≜ *p* < 0.0001.

**Table 1.** Current densities and gating properties of Ca_V_1 constructs expressed with and without α_2_δ-1 in GLT-C9 (Ca_V_1.1/α_2_δ-1-null) myotubes.

| **Constructs** | **N** | **n** | **I<sub>Ca</sub> (pA/pF)** | **P value** | **delta** | **V<sub>0.5</sub> (mV)** | **P value** | **delta** | **T<sub>activation</sub> (ms)** | **P value** | **fold change** |
| --- | --- | --- | --- | --- | --- | --- | --- | --- | --- | --- | --- |
| Cav1.1e | 4 | 10 | -8.9 ± 0.9 |  |  | 6.1 ± 1.9 |  |  | 6.0 ± 0.5 |  |  |
| Cav1.1e + $\alpha_2\delta$-1 | 4 | 10 | -6.9 ± 0.9 | 0.0717 | 2.0 | 7.6 ± 1.2 | 0.5039 | 1.5 | 20.0 ± 2.8 | 0.0007 | 0.3 |
| Cav1.2 | 9 | 10 | -8.4 ± 0.9 |  |  | 32.2 ± 2.4 |  |  | 4.5 ± 0.4 |  |  |
| Cav1.2 + $\alpha_2\delta$-1 | 3 | 5 | -12.7 ± 2.3 | 0.1087 | 4.4 | 9.4 ± 2.4 | <0.0001 | 22.8 | 1.5 ± 0.2 | <0.0001 | 3.0 |
| Cav1.1e_K162A | 3 | 9 | -6.8 ± 0.8 |  |  | 7.8 ± 2.6 |  |  | 2.9 ± 0.2 |  |  |
| Cav1.1e_K162A + $\alpha_2\delta$-1 | 4 | 9 | -5.0 ± 0.4 | 0.0426 | 1.9 | 3.1 ± 3.3 | 0.2581 | 4.7 | 1.8 ± 0.2 | 0.0167 | 0.6 |
| Cav1.1e_E90A | 3 | 9 | -3.2 ± 0.4 |  |  | 26.4 ± 1.0 |  |  | 4.3 ± 0.4 |  |  |
| Cav1.1e_E90A + $\alpha_2\delta$-1 | 3 | 9 | -2.5 ± 0.3 | 0.2073 | 0.7 | 16.0 ± 2.3 | 0.0014 | 10.4 | 2.8 ± 0.3 | 0.0069 | 0.7 |

### Molecular dynamics simulations reveal a direct molecular link between the α_2_δ-1 MIDAS domain and Ca_V_1.1 VSD I

The differential effects of α_2_δ-1 knockout on Ca_V_1.1 and Ca_V_1.2 activation kinetics indicates that α_2_δ-1 facilitates mechanisms regulating current kinetics that are intrinsic to the pore-forming channel subunits. Previous work identified VSD I as the prime regulator of activation kinetics of Ca_V_1 channels ^9–11^ and described interactions between VSD I and the functionally important MIDAS domain of α_2_δ-1 ^4,6^. Therefore, we hypothesized that direct interactions between α_2_δ-1 and VSD I of Ca_V_1.1 mediate the action of α_2_δ-1, which slows Ca_V_1.1’s activation kinetics. To identify the molecular details determining this functionally important interaction, we interrogated our structure model of Ca_V_1.1 in complex with α_2_δ-1, which is built on the cryo-EM structure of the calcium channel complex isolated from skeletal muscle ^4,7^ (**Fig. 3A**). In the activated state, critical residues of the α_2_δ-1 MIDAS domain formed hydrogen bonds with residues of a negatively charged cluster of amino acids located in the L1-2 loop of VSD I. Importantly, in α_2_δ-1 S263 and S265, the two serines of the canonical DxSxS MIDAS motif, interact with D78 of the Ca_V_1.1 L1-2_I_ loop (**Fig. 3B**). Notably, the structure further revealed that within the VSD I L1-2_I_ loop E76 and D77 interact with K162 in IS4. This is the first in a series of positively charged amino acids in the S4 helices considered as gating charges in the voltage-sensing process. Moreover, our recent study demonstrated that K162 plays an important role in determining the slow activation kinetics of Ca_V_1.1 ^12^. Thus, our structure model suggests that the negatively charged triplet E76/D77/D78 of the L1-2_I_ loop of VSD I forms the link connecting the α_2_δ-1 subunit with the outermost gating charge K0 (K162) of the IS4 helix. This could be the structural connection underlying the kinetics-regulating function of α_2_δ-1.

**Fig. 3.**
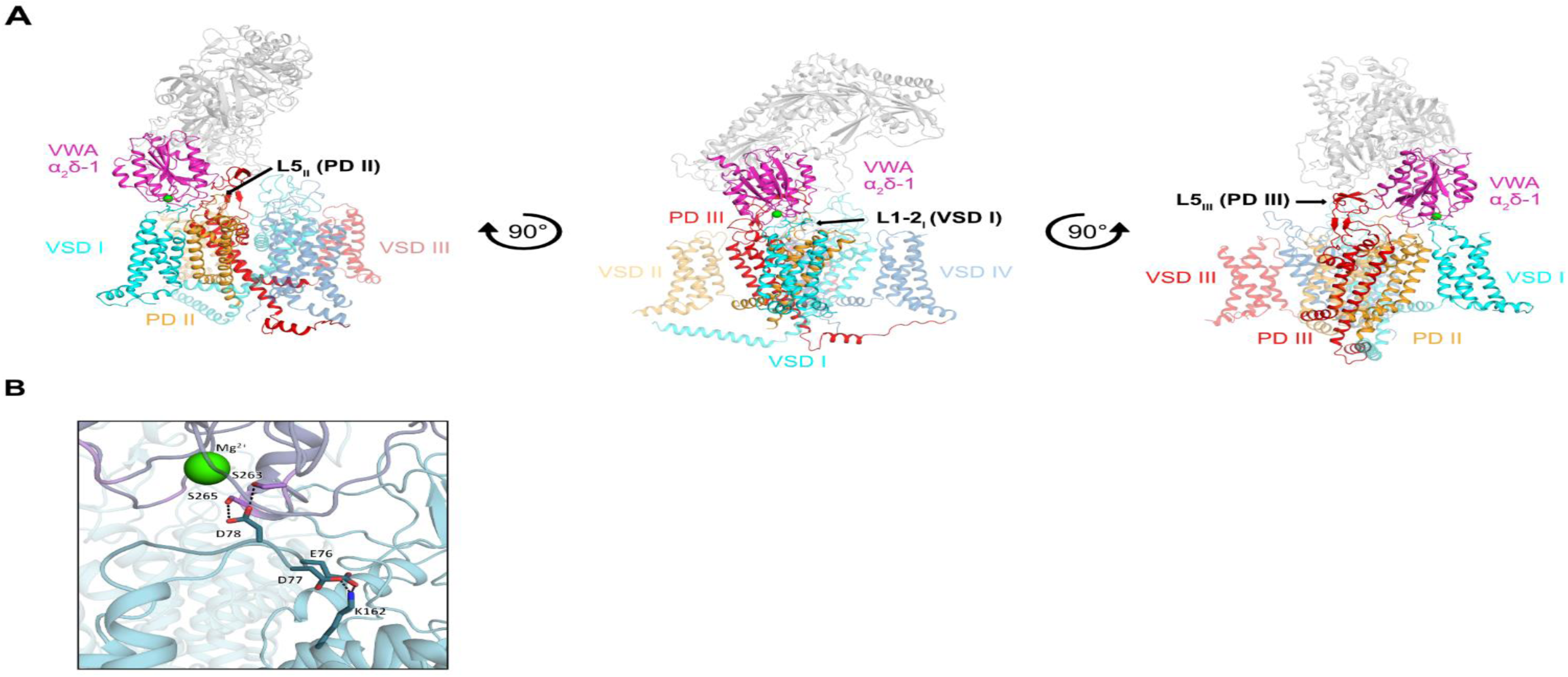
Structure prediction of the α_2_δ-1 – Ca_V_1.1 VSD I interaction site. **A**: Structure of the Ca_V_1.1/α_2_δ-1 complex (PDB code: 5GJV; ^4^) displaying the interaction sites between the two calcium channel subunits. Note that only one of the four VSDs (VSD I, cyan) interacts with α_2_δ-1, directly with the MIDAS site in the Von Willebrand A domain (VWA, magenta). Additional α_2_δ-1 interaction sites exist with the L5 loops of the pore-domains (PD) II and III. **B**: Close-up of the interaction site between the Ca_V_1.1 L1-2_I_ loop (cyan) and the MIDAS domain of α_2_δ-1 (magenta). D78 in the L1-2_I_ loop interacts with S265 and S263 of α_2_δ-1. The adjacent E76 and D77 in the L1-2_I_ loop both interact with K162 (K0) on the outer end of the IS4 helix.

### Alanine substitutions of the negatively charged amino acids in the L1-2_I_ loop accelerate the activation kinetics of Ca_V_1.1 and differentially affect α_2_δ-1 association with Ca_V_1.1

To examine the prediction of our structure model that the residues of the E76/D77/D78 cluster communicate the interaction between α_2_δ-1 and VSD I, we substituted the negatively charged glutamine and aspartates with alanines to interrupt their bilateral interactions with α_2_δ-1 and K162, respectively. As D78 in the L1-2_I_ loop formed the inter-subunit connection with S263 and S265 in α_2_δ-1, critical residues of the MIDAS motif, we examined the functional consequence of this interaction using a single amino acid substitution D78A. Since our structure indicated that E76 and D77 together form the intra-domain connection to K162, we initially examined their importance together in a double mutation E76A/D77A. Patch-clamp analysis in GLT myotubes (Ca_V_1.1-null, expressing endogenous α_2_δ-1) transfected with either Ca_V_1.1e_E76A/D77A or Ca_V_1.1e_D78A revealed the hypothesized acceleration of the activation kinetics (**Fig. 4D**). In both constructs, the time constants of the current activation were equally reduced by 2.5-fold relative to matched wildtype controls. This acceleration was similar to that of Ca_V_1.1e in the absence of α_2_δ-1 in GLT-C9 myotubes compared to the plus α_2_δ-1 condition (c.f. **Fig. 2G**). The current densities and voltage-dependence of activation were not different from controls, demonstrating the specificity of the effects on the regulation of activation kinetics. These, results support a specific role of the negatively charged cluster in L1-2_I_ in the functional connection of the α_2_δ-1 to Ca_V_1.1 VSD I.

**Fig. 4.**
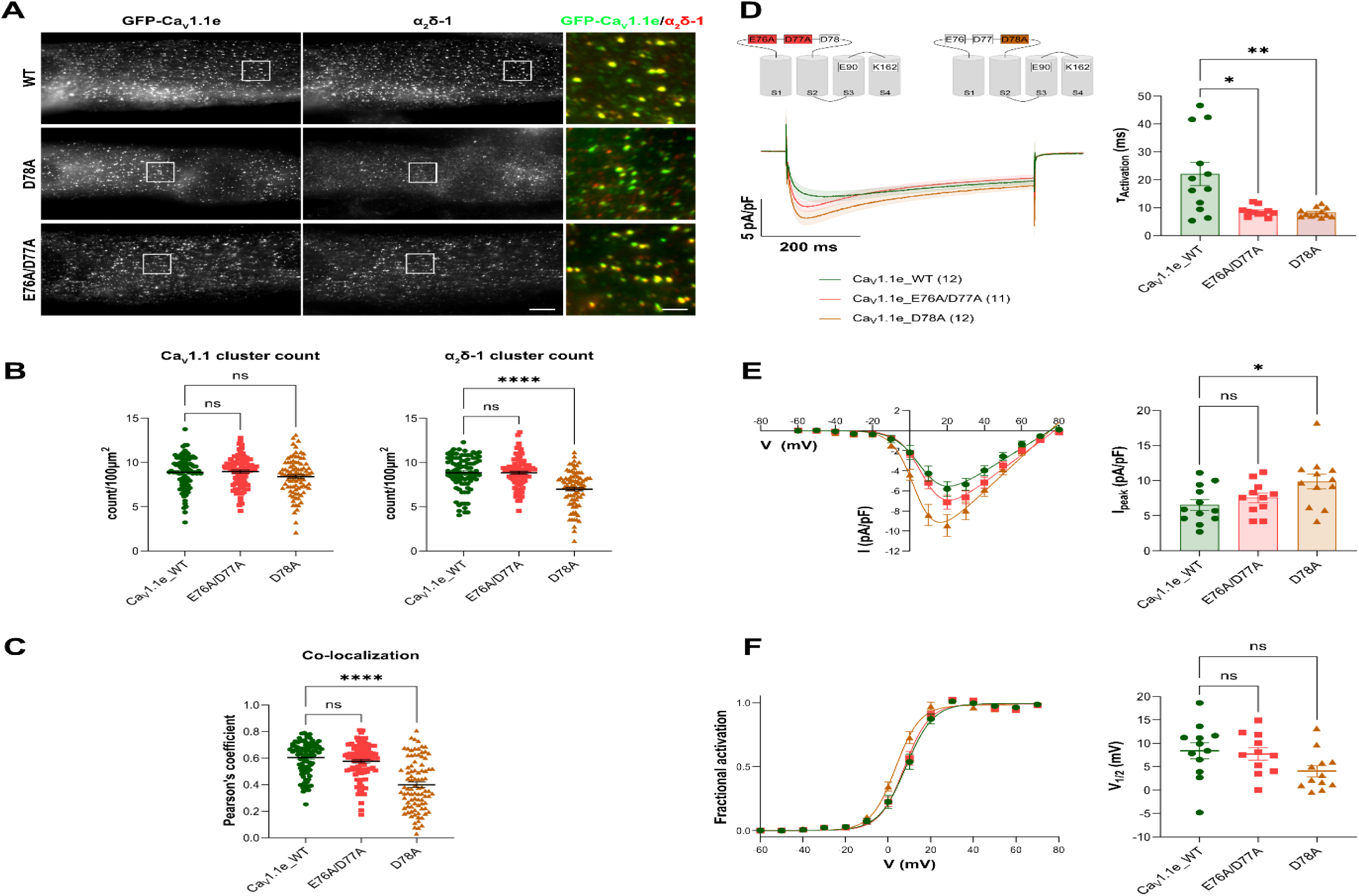
Alanine substitutions of negatively charged amino acids in L1-2_I_ differentially affect α_2_δ-1 association with Ca_V_1.1 and abolish slow calcium current activation. **A**: Double-immunofluorescence labeling of recombinant Ca_V_1.1 constructs and endogenous α_2_δ-1 demonstrates the co-clustering of wildtype and mutant constructs in triad junctions of GLT myotubes. **B,C**: Quantitative analysis demonstrates that the density of α_2_δ-1 clusters and the co-localization with Ca_V_1.1 are reduced in Ca_V_1.1e_D78A but not significantly altered in Ca_V_1.1e_E76A/D77A. Statistical tests: panel D, Kruskal-Wallis with Dunn’s multiple comparison; panel C, Welch’s ANOVA test with Dunnett’s T3 multiple comparison. **D-F**: Current properties recorded in GLT myotubes reconstituted with Ca_V_1.1_D78A and Ca_V_1.1_E76A/D77A mutant. Both mutations accelerate current activation equally by about 2.5-fold (**D**). Current density (**E**) and voltage-dependence of activation (**F**) are not altered. Statistical tests: panel D, Kruskal-Wallis with Dunn’s multiple comparison; panel E and F, One-way ANOVA with Dunnett’s multiple comparison. The data points are represented as mean ± SEM. ∗ ≜ *p* < 0.05, ∗∗ ≜ *p* < 0.01, ∗∗∗ ≜ *p* < 0.001, ∗∗∗∗ ≜ *p* < 0.0001.

Considering the interaction of D78 with α_2_δ-1 observed in the structure model (c.f. **Fig. 3B**) and the similarity of the kinetic effects in myotubes expressing the D78A mutation and when wildtype Ca_V_1.1e was expressed in the absence of α_2_δ-1 (cf. **Fig. 2G**), the D78A mutation might exert its effect simply by dissociating α_2_δ-1 from Ca_V_1.1. We examined this possibility by analyzing the co-localization of α_2_δ-1 with the D78A and E76A/D77A constructs expressed in GLT myotubes. Consistent with a role of D78 in anchoring α_2_δ-1 to Ca_V_1.1, α_2_δ-1 clustering with Ca_V_1.1e_D78A was significantly reduced to 79% and co-localization to 66% of that observed in matched wildtype control myotubes (**Fig. 4B,C**). In contrast, α_2_δ-1 clustering and co-localization of α_2_δ-1 with the Ca_V_1.1e_E76A/D77A double mutant immunofluorescence was not affected. Because mutation of D78 caused only a partial dissociation of α_2_δ-1 from Ca_V_1.1 and the combined mutation of E76 and D77 no detectable dissociation at all, we conclude that the functional effects of the alanine substitutions resulted from disrupting the transduction of the α_2_δ-1 effect to the speed control mechanism of VSD I, rather than from the physical dissociation of α_2_δ-1 from Ca_V_1.1.

### Combining mutations of E76A/D77A and D78A in a triple-mutation does not increase the kinetic effect above that of the separate mutations

If this interpretation is correct, and the D78 and E76/D77 residues function as the two arms connecting α_2_δ-1 with the speed-regulating mechanism of VSD I, combining the mutations should not result in an additive kinetic effect. To examine this prediction, we generated the triple-mutation Ca_V_1.1e_E76A/D77A/D78A and analyzed its current properties in GLT myotubes. Indeed, we observed a two-fold acceleration of activation kinetics in the Ca_V_1.1e_E76A/D77A/D78A triple-mutation relative to matched wildtype controls. This change was of approximately the same magnitude as that observed in the separate mutations shown above, and therefore does not indicate an additive effect on current kinetics when both the link to α_2_δ-1 (D78A) and the link to the IS4 helix (E76A/D77A) were severed simultaneously (**Fig. 5A-C**). The absence of an additive effect is consistent with the notion that both D78 and E76/D77 function as sequential components of the same signaling cascade.

**Fig. 5.**
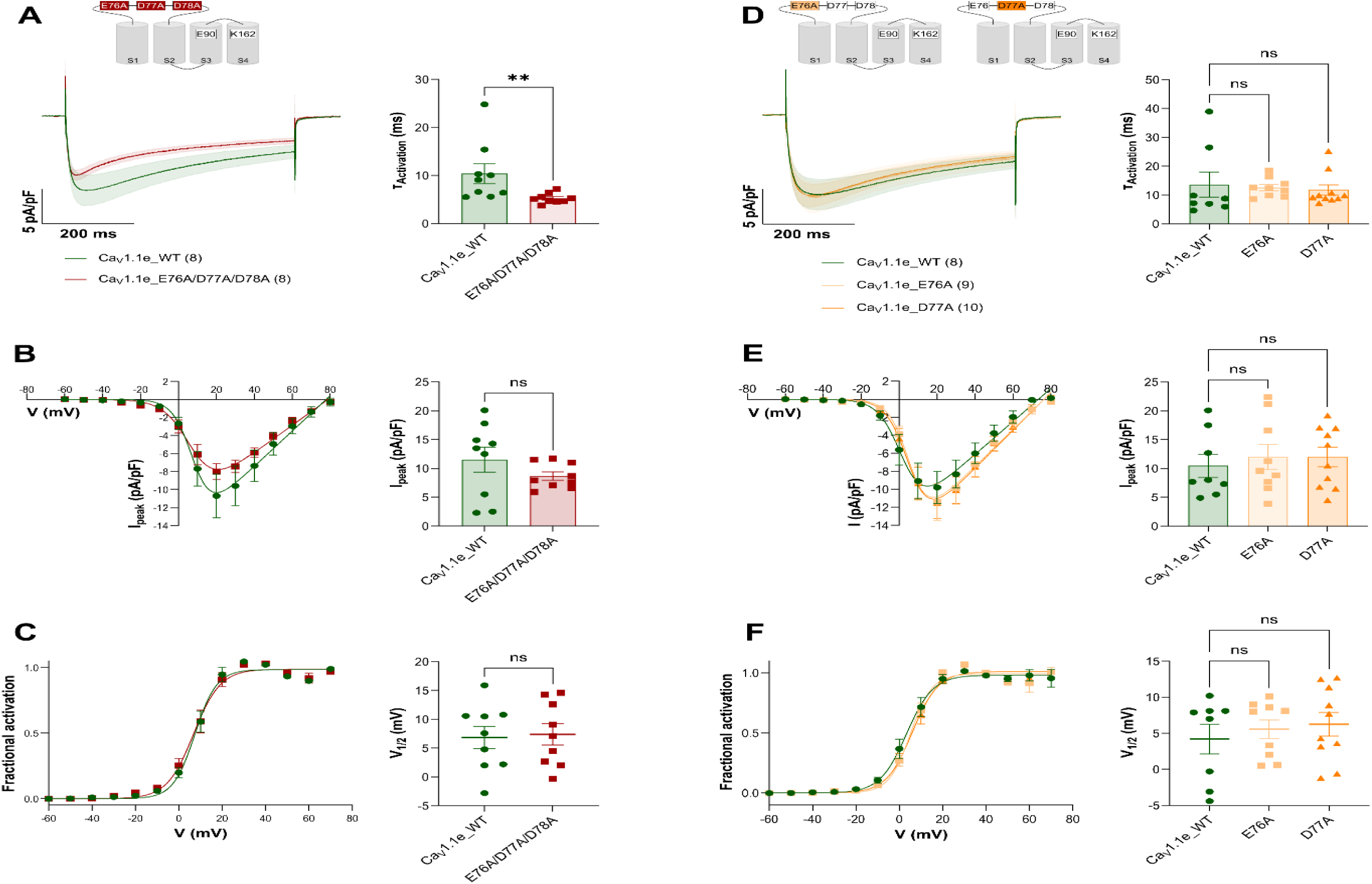
Individual and triple-mutations of E76, D77, and D78 in the L1-2_I_ loop of Ca_V_1.1 expressed in GLT myotubes. **A-C**: Current properties of Ca_V_1.1e_E76A/D77A/D78A triple-mutant compared to matched wildtype Ca_V_1.1e. Mean current traces and time constants of current activation (**A**) show a 2-fold acceleration of current kinetics on neutralization of all three negatively charged residues of the L1-2_I_ motif. Current density (**B**) and voltage-dependence of activation (**C**) were not altered. **D-F**: Current properties of individually mutated Ca_V_1.1e_ E76A and Ca_V_1.1e_D77A show no difference compared to wildtype Ca_V_1.1e. Current kinetics (**D**), current density (**E**) and voltage-dependence of activation (**F**) did not significantly differ between wildtype and mutant channels. The data points are represented as mean ± SEM. Statistical tests: panel A, Mann-Whitney test; panel B, Welch’s t-test; panel C, t-test; panel D-F, Kruskal-Wallis with Dunn’s multiple comparison. ∗ ≜ *p* < 0.05, ∗∗ ≜ *p* < 0.01, ∗∗∗ ≜ *p* < 0.001, ∗∗∗∗ ≜ *p* < 0.0001.

### Individual E76A and D77A substitutions are not sufficient to reproduce the effects of the double mutant

As mentioned above, the signaling arm connecting the negative amino acid cluster in the L1-2_I_ loop with K162 in IS4 actually consists of two potential binding partners, E76 and D77. Are both of these interactions necessary or is one or the other sufficient for transmitting the effects of α_2_δ-1 to VSD I? This question was examined using two individual mutations, Ca_V_1.1e_E76A and Ca_V_1.1e_D77A (**Fig. 5D-F)**. The electrophysiological analysis in GLT myotubes showed that neither one of these amino acid substitutions by itself altered the current kinetics of Ca_V_1.1e. This indicates that either one of the two possible connections of the negative amino acid cluster in L1-2_I_ to K162 is sufficient for transmitting the full modulatory effect of α_2_δ-1 to the speed control mechanism of VSD I.

### Kinetic effects of mutating two speed-controlling residues in VSD I (K162, E90) exceed effects of functional uncoupling of α_2_δ-1

The data shown up to this point indicate that the negatively charged amino acids in L1-2_I_ mediate the kinetic action of α_2_δ-1 on VSD I. Moreover, our structure model predicts direct interactions between residues E76/D77 and K162 in IS4. Consistent with a role in the regulation of current kinetics, recently, we reported that an alanine substitution of K162 strongly accelerated the activation kinetics of Ca_V_1.1 ^12^. K162 represents the outermost “gating charge” in the IS4 helix (also named K0), which is conserved in all high-voltage-activated Ca_V_ channels. While it is one of the positively charged residues located in three amino-acid intervals on the moving S4 helix, it remains on the extracellular side in the activated and resting states of VSD I and probably does not participate directly in the voltage-sensing process.

To examine whether K162 functions as the effector of the α_2_δ-1 action mediated by the L1-2_I_ EDD motif, we combined the alanine substitution of K162 (K162A) with the D78A mutation, to functionally dissociate VSD I from α_2_δ-1. If the only function of K162 is to link VSD I to α_2_δ-1, then this double mutant would be expected to yield the same accelerating effect as disruption of the L1-2_I_ link in the single mutation D78A, double mutation E76A/D77A or the triple mutation E76A/D77A/D78A. Alternatively, if K162 possesses an intrinsic function in regulating the kinetics of VSD I, an additive effect of the two mutations might be observed. As expected in either models, the double mutant Ca_V_1.1e_D78A/K162A caused an acceleration of activation kinetics (**Fig. 6A**). Current density and the voltage-dependence of activation were not altered (**Fig. 6B,C**), indicating its specific function in the regulation of the slow Ca_V_1.1 current kinetics. Notably, the time constant of current activation in Ca_V_1.1e_D78A/K162A expressing myotubes (τ 3.2 ms) was substantially faster than that of any of the L1-2_I_ EDD mutants (τ 5.8 – 8.5 ms; **Table 2**). This indicates that K162 possesses an intrinsic activity regulating activation kinetics, beyond that exerted by α_2_δ-1. Thus, defining K162 as part of the intrinsic speed control mechanism of Ca_V_1.1 VSD I.

**Fig. 6.**
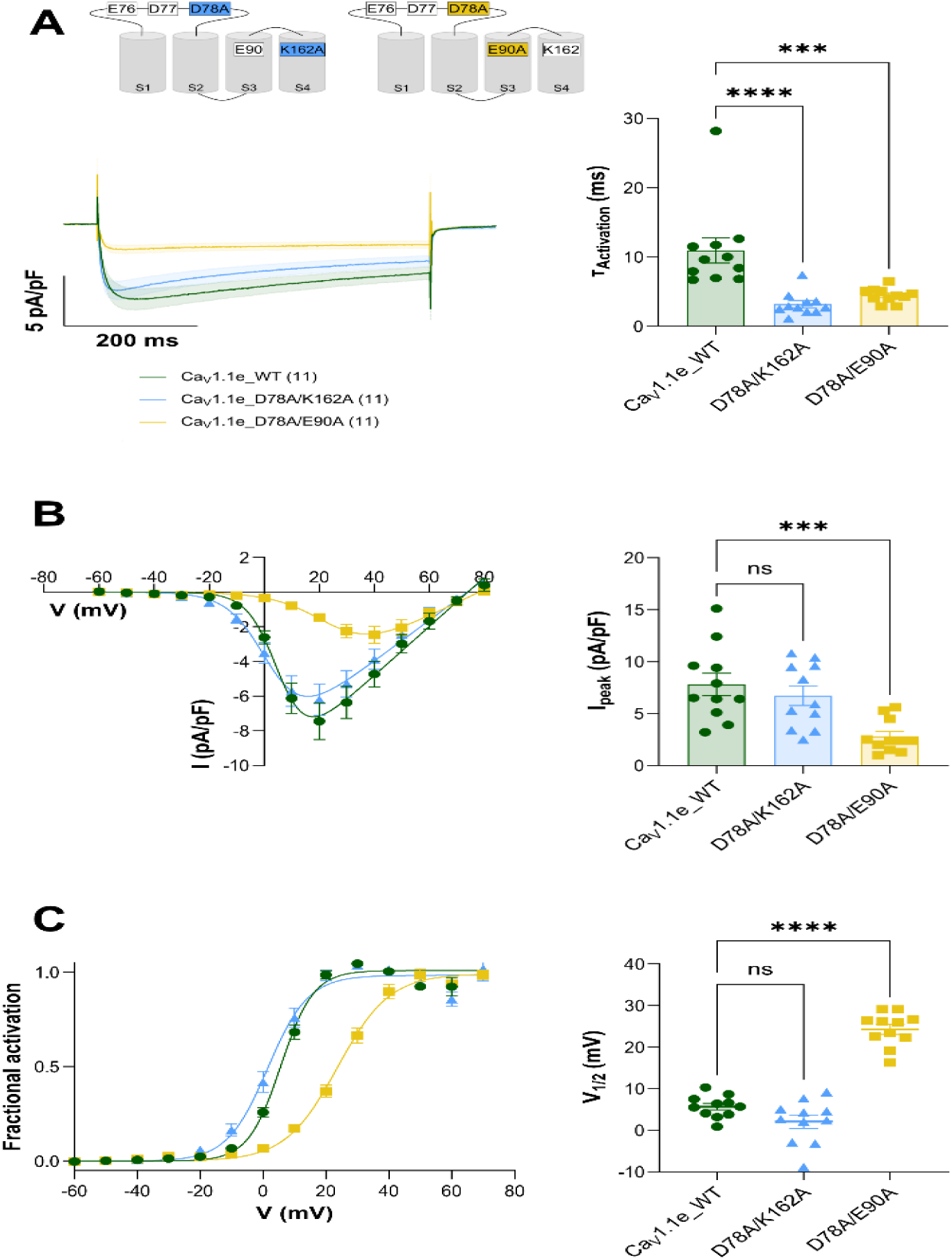
Acceleration of current activation by mutation of the outermost gating charge K162A and the countercharge E90A together with the L1-2_I_ mutation D78A. **A**: Mean calcium currents in GLT myotubes reconstituted with the double mutants Ca_V_1.1_D78A/K162A or Ca_V_1.1_D78A/E90A. Compared to wildtype controls current activation is accelerated in both double mutants. **B**: I/V curves show reduced current density for D78A/E90A. **C**: Activation curves and mean voltage of half-maximal activation shows a shift to less depolarizing potentials for D78A/E90A. The data points are represented as mean ± SEM. Statistical tests: panel A-C, One-way ANOVA with Dunnett’s multiple comparison. ∗ ≜ *p* < 0.05, ∗∗ ≜ *p* < 0.01, ∗∗∗ ≜ *p* < 0.001, ∗∗∗∗ ≜ *p* < 0.0001.

**Table 2.** Current densities and gating properties of Ca_V_1 constructs expressed in GLT (Ca_V_1.1-null) myotubes.

| Constructs | N | n | I<sub>Ca</sub> (pA/pF) | P value | delta | V<sub>0.5</sub> (mV) | P value | delta | T<sub>activation</sub> (ms) | P value | fold change |
| --- | --- | --- | --- | --- | --- | --- | --- | --- | --- | --- | --- |
| Ca<sub>v</sub>1.1e | 4 | 12 | -6.5 ± 0.8 |  |  | 8.4 ± 1.7 |  |  | 22.1 ± 4.1 |  |  |
| Ca<sub>v</sub>1.1e_E76A/D77A | 5 | 11 | -7.6 ± 0.7 | 0.6147 | 1.0 | 7.7 ± 1.3 | 0.9263 | 0.7 | 8.5 ± 0.5 | 0.0372 | 0.4 |
| Ca<sub>v</sub>1.1e_D78A | 5 | 12 | -9.9 ± 1.0 | 0.0156 | 3.4 | 4.1 ± 1.0 | 0.0695 | 4.3 | 8.5 ± 1.0 | 0.0088 | 0.4 |
| Ca<sub>v</sub>1.1e | 3 | 8 | -11.5 ± 2.2 |  |  | 6.8 ± 1.9 |  |  | 10.4 ± 2.1 |  |  |
| Ca<sub>v</sub>1.1e_E76A/D77A/D78A | 4 | 8 | -8.7 ± 0.7 | 0.2471 | 2.8 | 7.4 ± 1.9 | 0.8383 | 0.6 | 5.3 ± 0.3 | 0.0019 | 0.5 |
| Ca<sub>v</sub>1.1e | 3 | 8 | -10.5 ± 2.0 |  |  | 4.2 ± 2.1 |  |  | 13.6 ± 4.4 |  |  |
| Ca<sub>v</sub>1.1e_E76A | 4 | 9 | -12.0 ± 2.2 | >0.9999 | 1.5 | 5.6 ± 1.3 | >0.9999 | 1.4 | 12.6 ± 1.1 | 0.1435 | 0.9 |
| Ca<sub>v</sub>1.1e_D77A | 4 | 10 | -12.0 ± 1.7 | >0.9999 | 1.5 | 6.3 ± 1.0 | 0.7901 | 2.1 | 11.8 ± 1.0 | 0.7907 | 0.9 |
| Ca<sub>v</sub>1.1e | 4 | 11 | -7.8 ± 1.1 |  |  | 5.7 ± 0.8 |  |  | 11.0 ± 1.8 |  |  |
| Ca<sub>v</sub>1.1e_D78A/K162A | 3 | 11 | -6.7 ± 0.5 | 0.5888 | 1.1 | 2.1 ± 1.6 | 0.0878 | 3.6 | 3.2 ± 0.5 | <0.0001 | 0.3 |
| Ca<sub>v</sub>1.1e_D78A/E90A | 3 | 11 | -2.8 ± 0.5 | 0.0006 | 5.0 | 24.3 ± 1.2 | <0.0001 | 18.6 | 4.5 ± 0.3 | 0.0037 | 0.3 |

In the resting and intermediate states, K162 interacts with countercharges E87 and E90 in IS2 and with E140 at the extracellular end of IS3 (see **Fig. 3B**). Similar to K162A, alanine substitution of E90 has been shown to accelerate activation kinetics of Ca_V_1.1 ^11^. E90 represents a highly conserved countercharge in the IS2 helix, which forms consecutive interactions with S4 gating charges as they move up and down with the changing electrical potential. Because E90 is an important determinant of Ca_V_1.1’s slow activation ^11^, but, to our knowledge, is not directly structurally linked to the EDD motif, we wondered whether its function in regulating current kinetics also depends on the interaction with the α_2_δ-1 subunit via the L1-2_I_ EDD motif? To address this question, we combined the alanine substitution of E90 (E90A) with the D78A mutation, to functionally dissociate VSD I from α_2_δ-1. Consistent with previous results from the single E90A mutation ^11^, the Ca_V_1.1e_D78A/E90A double mutant caused an acceleration of current activation (**Fig. 6A**). Moreover, the current density was reduced and the voltage-dependence of activation was shifted to more positive potentials (**Fig. 6B,C**). Notably, as shown for the Ca_V_1.1e_D78A/K162A double mutant (above), the time constant of the Ca_V_1.1e_D78A/E90A double mutant (τ 4.5 ms) was faster than that of D78A alone (τ 8.5 ms; **Table 2**). Together, these experiments indicate that both K162 and E90 are components of an intrinsic speed control mechanism in VSD I, which can function independently of α_2_δ-1. Their fast activation time constants in the double mutants could either exhibit the intrinsic kinetic properties of K162 and E90 (thus masking any parallel effects of α_2_δ-1). Or the fast activation time constants could result from additive effects of the intrinsic K162 and E90 actions, plus that of α_2_δ-1. However, whether the VSD I intrinsic actions of K162 and E90 are entirely independent of α_2_δ-1 or modulated by it remains to be shown.

### Knockout of α_2_δ-1 reduces the effects on Ca_V_1.1 current kinetics of neutralizing mutations K162A and E90A

To address this possibility directly, we turned to the GLT-C9 cell line and expressed the individual alanine substitutions of the two intrinsic speed-controlling residues, K162A and E90A, with and without α_2_δ-1 (**Fig. 7**). Consistent with the data of the previous experiment, both Ca_V_1.1e_K162A and Ca_V_1.1e_E90A expressed in GLT-C9 cells without α_2_δ-1 showed substantially accelerated activation kinetics compared to wildtype Ca_V_1.1e expressed with or without α_2_δ-1 (**Fig, 7A,D; cf. Fig. 2A,G**). As expected, the time constants in the absence of α_2_δ-1 (K162A: τ 2.9 ms; E90A: τ 4.3 ms) were very similar to those recorded in GLT cells expressing the double mutants with the severed L1-2_I_ connection to α_2_δ-1 (D78A; **Table 2**). Unexpectedly, however, co-expression of α_2_δ-1 with either Ca_V_1.1e_K162A or Ca_V_1.1e_E90A further accelerated their activation kinetics by approximately two-fold (K162A + α_2_δ-1: τ 1.8 ms; E90A + α_2_δ-1: τ 2.8 ms). This effect of α_2_δ-1 in the two mutants clearly demonstrates that the intrinsic speed control mechanism in VSD I, represented by K162 and E90, is modulated by α_2_δ-1. The unexpected accelerating direction of this modulation indicates that the presence of the α_2_δ-1 subunit facilitates the accelerating effects of the K162A and E90A mutations, just as in wildtype Ca_V_1.1 it facilitates the intrinsic speed control mechanism in the opposite direction, slowing activation kinetics.

**Fig. 7.**
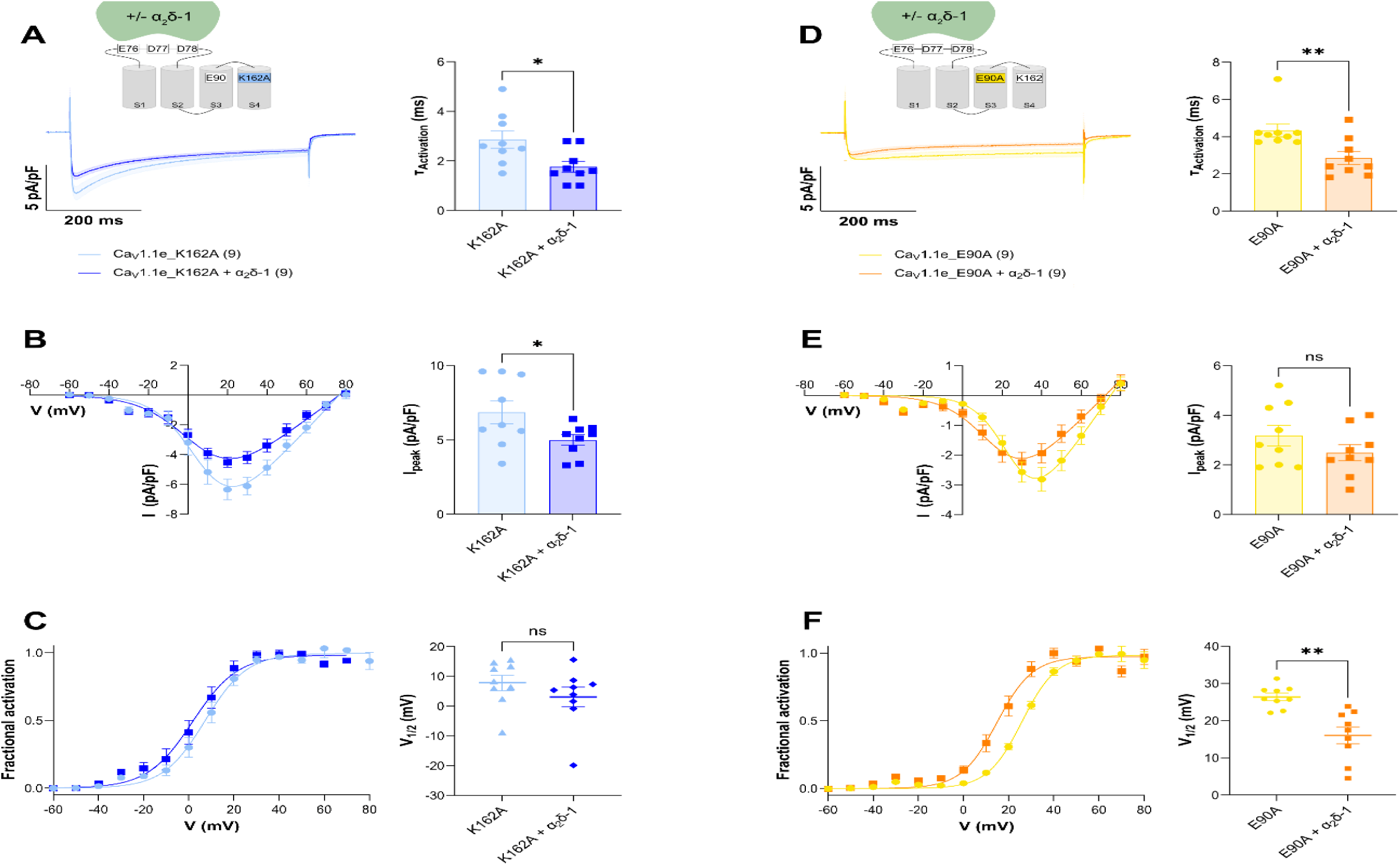
Further current acceleration of charge-neutralizing mutations K162A and E90A by co-expression of α_2_δ-1 in GLT-C9 myotubes. **A,D**: Mean calcium currents in GLT-C9 myotubes reconstituted with Ca_V_1.1_K162A or Ca_V_1.1_E90A with and without α_2_δ-1. In the presence of α_2_δ-1 current kinetics are further accelerated in both mutants. **B,E**: I/V curves and mean current densities. **C,F**: Activation curves and mean voltage of half-maximal activation shows a shift to less depolarizing potentials for E90A with α_2_δ-1. The data points are represented as mean ± SEM. Statistical tests: panel A, D and E, Mann-Whitney test; panel B and E, t-test; panel C, Welch’s t-test. ∗ ≜ *p* < 0.05, ∗∗ ≜ *p* < 0.01, ∗∗∗ ≜ *p* < 0.001, ∗∗∗∗ ≜ *p* < 0.0001.

### Association of α_2_δ-1 stabilizes the L1-2_I_ loop and refines the interactions between the S4 gating charges and countercharges

The results of our mutagenesis experiments suggest that α_2_δ-1 modulates the intrinsic properties of VSD I through the EDD motif in the L1-2_I_ loop. To examine the underlying molecular mechanism, we generated structure models of Ca_V_1.1 VSD I with and without α_2_δ-1 in the activated, intermediate and resting states (**Fig. 8**). Our MD simulations of the state transitions in response to an applied electric field of Ca_V_1.1 without α_2_δ-1 showed three clearly discernable states: activated, intermediate and resting ^7^. Here we modeled each of these states in complex with the α_2_δ-1 subunit (**Fig. 8A**). As S4 moves from the resting down-to the activated up-position the IS4 gating charges sequentially interact with the countercharges of the CTC below the hydrophobic constriction site (HCS) and with those of the ENC above the HCS. Previous mutagenesis studies suggested that this repetitive breaking and formation of ionic interactions slowed the state transitions of VSD I and thus activation kinetics of Ca_V_1.1 ^11,12^. In the intermediate state K162 reaches up to the L1-2_I_ loop, forming ion-bonds with E76 and D77. This interaction is maintained in the activated state.

**Fig. 8.**
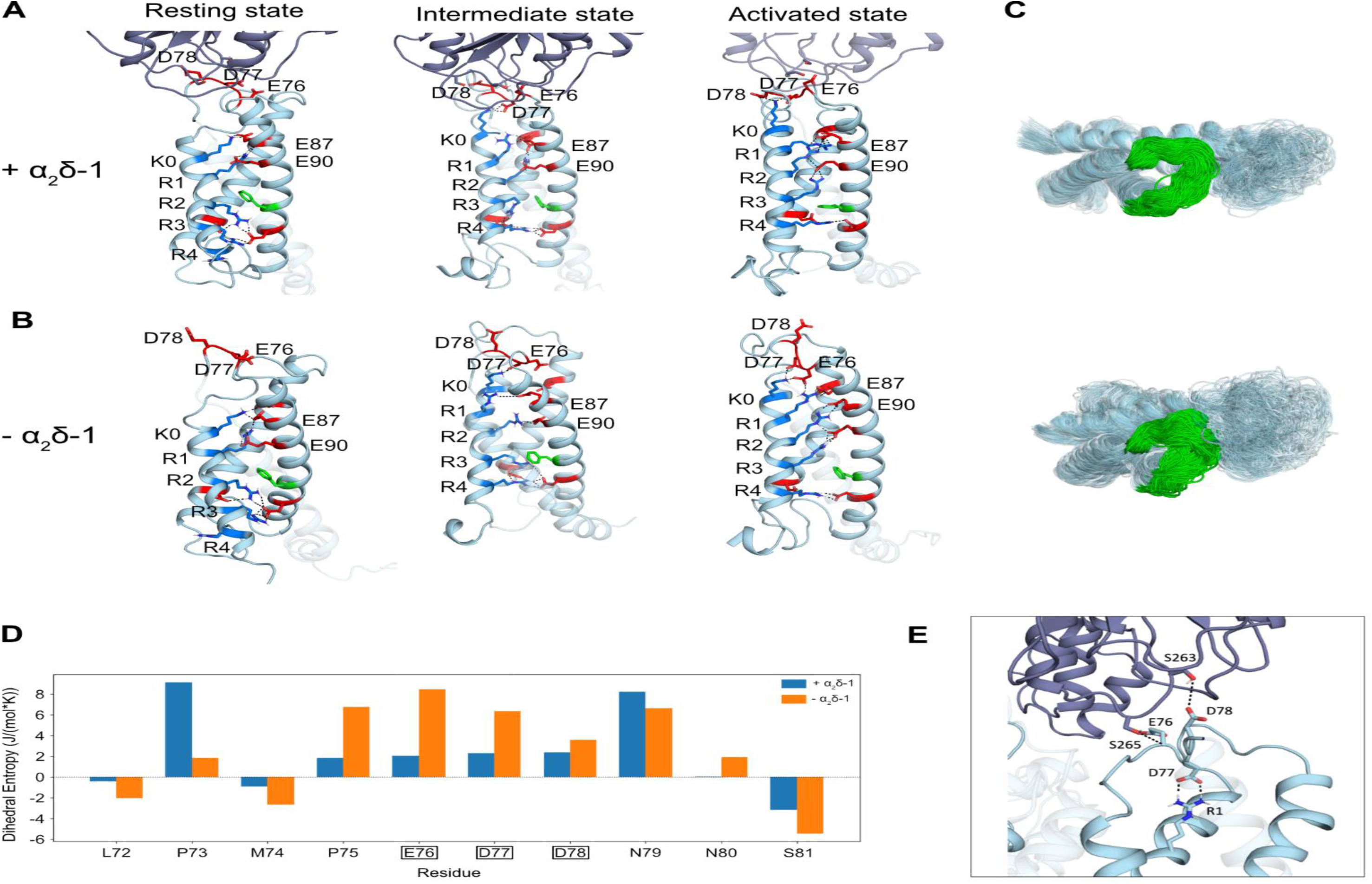
Molecular modeling of the state-transition of Ca_V_1.1 VSD I with and without α_2_δ-1. A: Structure of the Ca_V_1.1 VSD I (cyan) in complex with α_2_δ-1 (purple) in the resting, intermediate and activated state. **B:** Structure of the Ca_V_1.1 VSD I (cyan) without α_2_δ-1 in the resting, intermediate and activated state. S4 gating charges are shown in blue, countercharges in red and the phenylalanine of the HCS in green. Note, how in the absence of α_2_δ-1 the L1-2_I_ loop is flipped down and the K162 – E76/D77 interactions relaxed, the S2 helix is bent outward and the gating charge – countercharge interactions are relaxed. **C**: Overlay of cluster structures of VSD I (top view) with L1-2_I_ shown in green (104 clusters with α_2_δ-1, 303 clusters without α_2_δ-1, distance cut-off 2 Å). Without α_2_δ-1 the structures are less compressed, indicative of increased flexibility of the loop as well as the transmembrane helices. **D**: Dihedral entropies for all L1-2_I_ loop residues with (blue) and without (orange) α_2_δ-1. Without α_2_δ-1 the dihedral entropy of the residues forming the interaction site between VSD I and α_2_δ-1 (E76, D77 and D78) is higher, indicating an increased local flexibility. **E**: Model of the α_2_δ-1 – L1-2_I_ interface with a K162A substitution, showing that in the activated state R1 compensates for the lacking interaction with D77.

While this sequence of interactions between the S4 gating charges and the countercharges is seen with and without α_2_δ-1, the structural details of these interactions are greatly different in the two conditions. For example, with α_2_δ-1 the side chain of K0 points straight upward to E76/D77, whereas without α_2_δ-1 the L1-2_I_ loop flips downward, allowing interactions with K162 (K0) and R165 (R1). Apparently, the absence of α_2_δ-1 causes a greater flexibility of the L1-2_I_ loop as evidenced by significantly increased dihedral entropies of the L1-2_I_ loop residues (**Fig. 8 C,D**). Importantly, the lacking stabilization of the L1-2_I_ loop has striking consequences on the structure and position of S4 relative to the other helices of VSD I. Without the up-ward pull of the K0-E76/D77 interaction, in the absence of α_2_δ-1 the S4 helix seems to relax and consequently the S2 helix bends between its interaction sites on either sides of the HCS. This, in turn, appears to alter the interactions between the gating charges and the countercharges. Although, the specific consequences of each one of these changes is not known, the sum of these structural changes of the intrinsic activation mechanism of Ca_V_1.1 VSD I can readily explain the acceleration of activation kinetics observed in the absence of α_2_δ-1 or when the link to α_2_δ-1 is severed by one of the tested mutations.

Finally, we modeled the structure of VSD I of the K162A mutation in complex with α_2_δ-1 (**Fig. 8E**). In the absence of K162 the next gating charge down, R1, forms strong interactions with D77 in the L1-2_I_ loop. Thus, as R1 replaces K0 in this mutation, one of the steps in the up-ward state transitions is skipped, explaining the accelerated activation kinetics of K162A.

## Discussion

What is the role of the auxiliary α_2_δ-1 in the function of L-type calcium channels in muscle cells and how does α_2_δ-1 exert its function on the pore-forming α_1_ subunits? The results presented here indicate that α_2_δ-1 facilitates the intrinsic activation kinetics of Ca_V_1.1 and Ca_V_1.2. Moreover, our mutational analysis identified a contiguous string of ionic interactions between charged amino acids connecting the extracellular α_2_δ-1 subunit with the speed controlling mechanism in the first VSD of the Ca_V_1.1 L-type calcium channel. In Ca_V_1.1, alanine substitution of each of the involved amino acids caused an acceleration of current activation, similar to that recorded when α_2_δ-1 was eliminated from the Ca_V_1.1 channel complex (**Fig.9B**). At the core of this intramolecular signaling pathway is a cluster of three negatively charged amino acids (E76, D77, D78) located in the extracellular L1-2_I_ loop of VSD I, which serves as two-armed connector between the α_2_δ-1 subunit and the cytoplasmic end of the IS4 helix. Hydrogen bonds between the outwardly oriented D78 and the critical S263 and S265 in the MIDAS domain of α_2_δ-1 functionally connects the auxiliary subunit to the pore-forming channel subunit. Hydrogen bonds between one or both of the inwardly oriented E76 and D77 and the outermost gating charge K162 extend this connection to the IS4 helix **(Fig. 9A**). Sequential interactions between the IS4 gating charges, including K162, and the IS2 countercharges of the extracellular negative cluster including E90, appear to slow down the VSD I state transitions from the resting to the activated state in an α_2_δ-1-dependent manner.

**Fig. 9.**
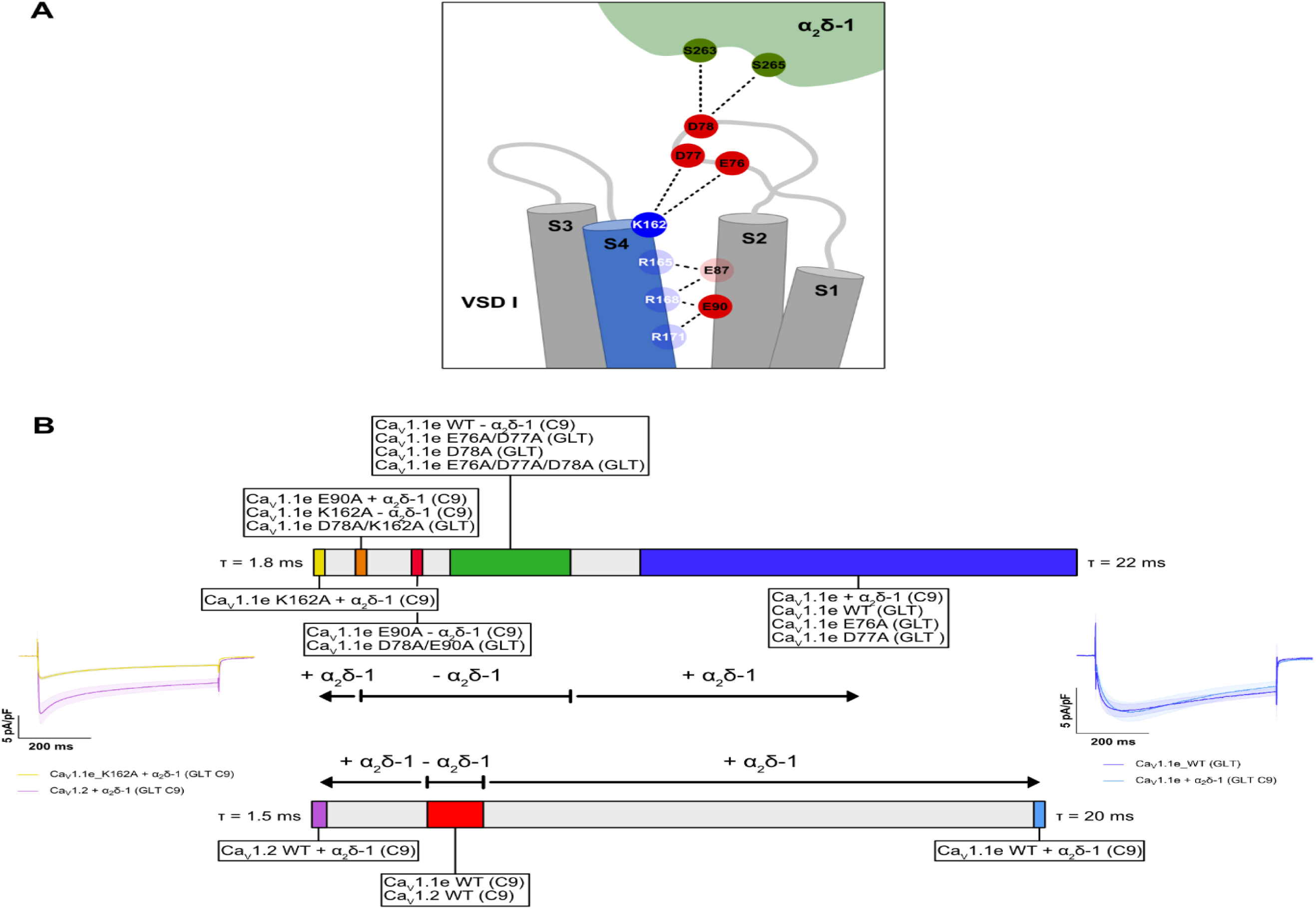
Schematic presentation of the α_2_δ-1 signaling cascade in Ca_V_1.1 VSD I and the comparative activation kinetics of the various tested channel constructs and subunit combinations. **A**: S263 and S265 of the α_2_δ-1 MIDAS motif interact with D78 in the extracellular L1-2_I_ loop of VSD I. The adjacent E76 and D77 interact with the outermost gating charge K162 of IS4, which in turn interacts with the countercharge E90 in IS2. Knockout of α_2_δ-1 and alanine mutations of all of these amino acids result in different degrees of acceleration of activation kinetics in Ca_V_1.1 expressed in skeletal muscle cells. **B**: Wildtype Ca_V_1.1e in combination with α_2_δ-1 activate with slow kinetics (blue range, τ: 10.4 – 22.1 ms). Knockout of α_2_δ-1 or functionally uncoupling of α_2_δ-1 by mutations in the L1-2_I_ EDD motif result in moderately accelerated activation kinetics (green range, τ: 5.3 – 8.5 ms). Mutating components of the intrinsic speed control mechanism results in even faster activation kinetics, which are further accelerated when α_2_δ-1 is present and functionally coupled to VSD I (K162A, orange, τ: 2.9 – 3,2 ms ◊ K162A + α_2_δ-1, yellow, τ: 1.8 ms; E90A, red, τ: 4.3 – 4.5 ms → E90A + α_2_δ-1, orange, τ: 2.8 ms). Therefore, the function of α_2_δ-1 is to make the VSD I with an intact speed control mechanism slow and that with a mutated speed control mechanism fast. This action is reminiscent of α_2_δ-1’s differential action of slowing the activation of Ca_V_1.1 and accelerating the activation of Ca_V_1.2.

Considering this arrangement, it is tempting to envision the connection to α_2_δ-1 functioning as a mere brake on the movement of IS4 that slows down its transitions from the resting into the activated state. However, our data do not support such a mechanism. Instead, α_2_δ-1 appears to facilitate the function of the intrinsic speed control mechanism within VSD I. First, this is suggested by our observation that the kinetic regulation of Ca_V_1.1 and Ca_V_1.2 by α_2_δ-1 acts in opposite directions. While a brake on the movement of IS4 could explain the decelerating action of α_2_δ-1 on Ca_V_1.1, it is inconsistent with the accelerating action of α_2_δ-1 on Ca_V_1.2. Secondly, we observed a similar accelerating action of α_2_δ-1 in the Ca_V_1.1 K162A and E90A mutants, which make Ca_V_1.1 VSD I behave in a Ca_V_1.2-like manner. Thus, the intrinsic gating charge – countercharge interactions in VSD I determine whether a channel activates slowly or rapidly. In both cases α_2_δ-1 linked to VSD I via L1-2_I_ EDD motif accentuates these divergent kinetic properties.

### What are the specific functions of the charged residues along the α_2_δ-1 – VSD I signaling path?

The data presented here suggest that the third residue of the EDD motif in the L1-2_I_ loop, D78, constitutes the major link to the MIDAS domain of α_2_δ-1. This is indicated by our structure model based on the cryo-EM structure of the skeletal muscle Ca_V_1.1 channel complex ^4,7^ and strongly supported by our immunolabeling experiments and functional analyses. Mutation of D78 reduces the incorporation of α_2_δ-1 into the channel complex at the triads and causes the same degree of acceleration of current kinetics as the knockout of α_2_δ-1. Moreover, a function of L1-2_I_ D78 as the primary link to α_2_δ-1 is consistent with a previous mutagenesis study that showed the critical importance of the corresponding residue in Ca_V_1.2 for the association and the functional effects of α_2_δ-1 to that L-type channel isoform^6^.

Our structure model indicates that in α_2_δ-1, both S263 and S265 of the canonical DxSxS MIDAS motif act as interaction partners of D78 in Ca_V_1.1. Mutations of this motif have previously been shown to disrupt α_2_δ-1 association to Ca_V_ α_1_ subunits and the effects of α_2_δ-1 on membrane expression and gating properties of calcium channels ^5,17^. Notably, our immunofluorescence analysis demonstrates that mutation of D78 causes only a partial reduction of α_2_δ-1 co-clustering, indicating that additional interactions substantially contribute to the association of the two channel subunits in skeletal muscle triads. This could be the interaction sites between α_2_δ-1 and the pore domains of repeats II and III of Ca_V_1.1 identified in the cryo-EM structure ^4^. Importantly however, our results indicate that the functional effects of α_2_δ-1 on Ca_V_1.1 current kinetics are completely lost when D78 is mutated, indicating that the interaction between the α_2_δ-1 MIDAS motif and D78 in VSD I is essential for the effects of α_2_δ-1 on Ca_V_1.1 channel properties. This further indicates that α_2_δ-1 modulates the channel gating properties at the level of the VSD and not at the pore domain.

Slow activation kinetics of Ca_V_1.1 were equally abolished when the two inward-pointing residues of the L1-2_I_ EDD motif (E76 and D77) were mutated simultaneously. The magnitude of this effect was identical to that of the D78A mutation. Moreover, when all three residues of the EDD motif were neutralized in a triple mutation, the effects were not additive, consistent with the notion that both mutations disrupted different parts of the same process. Immunofluorescence analysis demonstrated that, in contrast to D78A, the E76A/D77A double mutation did not substantially reduce co-clustering of α_2_δ-1 and Ca_V_1.1. From this observation, we conclude that the functional effects of mutations in the L1-2_I_ EDD motif were not simply caused by the physical dissociation of α_2_δ-1 from the channel, but by the disruption of the signaling pathway downstream of α_2_δ-1. Specifically, our structure analysis suggests ionic interactions of both negatively charged amino acids with the outermost IS4 gating charge K162. Our data further show that interactions of K162 with either E76 or D77 are sufficient for mediating the α_2_δ-1 effects; because single E76A or D77A mutations were not able to accelerate the activation kinetics.

Consistent with its pivotal role in the regulation of activation kinetics, alanine substitutions of K162 also accelerated Ca_V_1.1 activation kinetics ^12^. The interactions of E76 and D77 with K162 indirectly link α_2_δ-1 to the voltage-sensitive S4 helix of VSD I. This positively charged transmembrane helix is pulled inward by the negative resting potential and moves outward upon depolarization. The speed of VSD I state transitions from the resting to the activated state determines the activation kinetics of Ca_V_1.1 and critically depends on the interactions between the IS4 gating charges and countercharges in the adjacent helices ^11,12^. Unlike the interaction of R1 with IS2 countercharges E87 and E90, which stabilize the activated state, the interactions of K162 with E76 and D77 probably do not stabilize the activated state. The voltage-dependence of activation is not affected by alanine substitution of K162, E76 or D77. Accordingly, in Ca_V_1.1 voltage-dependence of activation is not modulated by α_2_δ-1.

However, the effect of these mutations on activation kinetics can be explained by the interactions of K162 in the resting and intermediate states. Our MD simulations indicate that before K162 reaches up to form the interactions with E76 and D77 in the L1-2_I_ loop it sequentially interacts with E90 and E87 positioned at the extracellular end of the IS2 helix. Repetitive formation and braking of these ionic bonds may explain the critical role of K162 in slowing down activation kinetics of VSD I. Conversely, mutations of K162 or the countercharge E90 each remove one step in this process. Consequently, the respective ion-bond partners will encounter one less energy barrier to overcome on their way up into the activated state. This speed-limiting mechanism also operates in the absence of α_2_δ-1 or when the L1-2_I_ link to α_2_δ-1 is severed by alanine substitutions, but the efficacy of this mechanism depends on its interaction with α_2_δ-1. Therefore, the role of K162 is two-fold: First, it contributes actively to the intrinsic speed control mechanism of VSD I, and secondly, it mediates the modulatory effects of α_2_δ-1 onto this mechanism.

In the transition of the IS4 helix from the resting to the activated state, K162 exchanges its interaction partners from E87/E90 to E76/D77. This breaking and formation of ionic bonds likely contributes to the slow activation of VSD I and is dependent on binding of the α_2_δ-1 MIDAS domain to the EDD motif in the L1-2_I_ loop. Our structure models indicate that in the absence of α_2_δ-1 the L1-2_I_ loop becomes much more flexible and its interactions are less defined (i.e. relatively short-lived). When the functional link to α_2_δ-1 is severed by mutating D78, the current-accelerating effect of K162A is reduced. Apparently, the accelerating effects of K162A and D78A are not additive. Instead, the very fast activation of K162A depends on the interaction of the L1-2_I_ loop with α_2_δ-1, just as the very slow activation of the wildtype Ca_V_1.1 depends on the α_2_δ-1 subunit. A possible explanation of this result comes from our observation that in the absence of K162, the next gating charge down, R1, interacts with E76/D77, possibly compensating for the loss of the K162 interaction. Apparently, also this compensatory interaction depends on the stability of the L1-2_I_ loop imparted by its interaction with α_2_δ-1. Likewise, the reduced kinetic effect observed in the E90A-D78A double mutant, can be explained by the weakened interaction of K162 with the unstable EDD motif uncoupled from α_2_δ-1. Together, these results demonstrate that both E90 and K162 are important components of the VSD I-intrinsic speed control mechanism, the function of which depends on the interaction of an outer gating charge (K0 in wildtype, R1 in the K162A mutant) with the EDD motif. In turn, this interaction requires the stabilization of the L1-2_I_ loop by α_2_δ-1.

### Role of the L1-2_I_ link in VSD I in mediating the differential effects of α_2_δ-1 on Ca_V_1.1 and Ca_V_1.2 calcium channels

Consistent with the results of previous studies from our group ^1–3^, the data obtained here in reconstituted Ca_V_1.1/α_2_δ-1 double-knockout myotubes clearly demonstrate that in the context of differentiated muscle cells the primary function of the α_2_δ-1 subunit is in regulating the activation kinetics of L-type calcium channels. Specifically, co-expression of α_2_δ-1 with the skeletal muscle Ca_V_1.1 isoform further slowed this channeĺs activation kinetics, and co-expression of α_2_δ-1 with the cardiac/neuronal Ca_V_1.2 isoform further accelerates that channeĺs activation kinetics. With Ca_V_1.2, we additionally observed a shift of the voltage-dependence of activation to more depolarizing potentials. This is in line with similar right-shifts reported in Ca_V_1.2/α_2_δ-1 co-expression studies in heterologous cells, in siRNA knock-down of α_2_δ-1 in myotubes, and in whole-animal knockout of α_2_δ-1 in mice ^2,6,18^. Together, the opposite kinetic effects and the differential effects on voltage-dependence of activation exerted by α_2_δ-1 on Ca_V_1.1 and Ca_V_1.2 expressed in myotubes corroborate the notion that the specificity of α_2_δ-1 effects is determined intrinsically in the α_1_ subunit, not by α_2_δ-1.

For Ca_V_1.1 a functional interaction between α_2_δ-1 and VSD I controlling current kinetics is consistent with the dominating role of this VSD in controlling channel gating ^9–12,19,20^. However, for Ca_V_1.2 voltage-clamp fluorometry studies suggested that VSDs II and III, rather than VSD I, dominate channel gating ^21^. Nevertheless, co-expression of Ca_V_1.2 with α_2_δ-1 increased the coupling efficiency to the pore of VSD I, II and III ^8^. Considering the distinct pore coupling of VSDs in Ca_V_1.1 ^19,22^ and Ca_V_1.2 ^21^ and the enhancing effects of α_2_δ-1 ^8^, it is conceivable that association of α_2_δ-1 alters the relative contribution to pore opening of VSDs with distinct kinetic properties. In Ca_V_1.1 + α_2_δ-1 the intrinsically slow VSD I dominates gating. In the absence of α_2_δ-1, its contribution to pore opening could be reduced and the relative contributions of the faster VSDs would be increased, thus causing the observed acceleration of activation kinetics. Conversely, in Ca_V_1.2 the energetic contribution of VSDs II and III to pore opening in channels lacking α_2_δ-1 is greatly reduced ^8^. This would increase the relative contribution of VSD I and thus cause the deceleration of current activation observed here.

While this model provides an attractive explanation of our current findings, it is in conflict with several lines of experimental evidence. First, even in the voltage-clamp fluorometry study of Ca_V_1.2 association of α_2_δ-1 caused a striking acceleration of the activation kinetics of VSD I (see Savalli et al., 2016, Fig. 3) ^8^, indicating a direct effect of α_2_δ-1 on VSD I properties. Also in Ca_V_1.2 the association of α_2_δ-1 via the EDD motif in VSD I is critical for the α_2_δ-1 regulation of the voltage-dependence of activation ^6^. Moreover, the activation kinetics of Ca_V_1.2 can be slowed down by swapping VSD I sequences with Ca_V_1.1 ^9^. Together, these functional studies support an important role of VSD I in determining gating properties and its modulation by α_2_δ-1 in both, Ca_V_1.1 and Ca_V_1.2. Furthermore, we consider it highly likely that the L1-2_I_ EDD motif transmits the modulatory effects of α_2_δ-1 in both Ca_V_1 channels and possibly throughout the entire family of high-voltage-activated Ca_V_ channels, in all of which the EDD motif is conserved.

The full conservation between Ca_V_1.1 and Ca_V_1.2 of the EDD motif and the downstream actuators K162 and E90 raises the question as to what causes the opposite effects of α_2_δ-1. Using a chimera approach, the laboratory of Kurt Beam and our own group has demonstrated that the kinetic properties of Ca_V_1.1 and Ca_V_1.2 can be exchanged by swapping of the IS3 helix and the adjacent extracellular IS3-S4 linker of the first VSD ^9,10^. Interestingly, all residues examined in the present study reside outside of these sequences (E76, D77, D78 in L1-2_I_, E90 in IS2 and K162 in IS4). Therefore, they represent a common mechanism of speed control in Ca_V_1.1 and Ca_V_1.2 and of its modulation by α_2_δ-1. Evidently, the sequence differences that render Ca_V_1.1 slowly activating and Ca_V_1.2 rapidly activating reside in the non-conserved amino acids of the interspersed IS3 helix and IS3-S4 loop. Differences in these sequences might cause changes in the functional interactions of the conserved players involved in the control of activation kinetics, including K162 and E90. This intrinsic speed control mechanism in VSD I is under the control of at least two supplementary structures: The intrinsic, isoform-specific features of the IS3 helix and the IS3-S4 loop, which determine the slow and fast activation of Ca_V_1.1 and Ca_V_1.2, respectively. And by the α_2_δ-1 subunit, which facilitates the expression of the distinct kinetic property via a common mechanism. The EDD motif in L1-2_I_ represents this mechanism, forming a two-armed connection between α_2_δ-1 and the intrinsic speed control mechanism in VSD I.

## Methods

### Plasmids

Cloning procedures for GFP-Ca_V_1.1e (Genbank NM_001101720) and GFP-Ca_V_1.1e-K162A were previously described ^12,23^. Sanger sequencing (Eurofins Genomics) verified the integrity of the newly generated plasmid.

GFP-Ca_V_1.1e-E76A, -D77A, -D78A, -E76A/D77A, -E76A/D77A/D78A, -E90A, -D78A/K162A, -D78A/E90A: To generate the desired constructs, each mutation was introduced into GFP-Ca_V_1.1e by splicing by overlap extension (SOE) PCR. First, the cDNA sequence of rabbit Ca_V_1.1 (nt 1–1006) was amplified in separate PCR reactions using GFP-Ca_V_1.1e as the template, employing overlapping primers that introduced the mutations. The two resulting PCR products were then used as templates for a PCR reaction with flanking primers to connect the nucleotide sequences. They were then combined in a subsequent reaction using flanking primers to connect the nucleotide sequences. The final fragment was digested with SalI and EcoRI and ligated into the corresponding sites of GFP-Ca_V_1.1e. The flanking primers used for all construct were: SalI-F: 5’-cgaaaagagagaccacat-3’ and EcoRI-R: 5’-cgtgatccagctcatgta-3.

The primers introducing the mutations were:

E76A, fw 5’-gcccatgcccgcggatgacaacaactccctgaacct-3’, rev 5’-tgttgtcatccgcgggcatgggcaggtacacggcca-3’.

D77A, fw 5’-catgcccgaggctgacaacaactccctgaacctggg -3’, rev 5’-agttgttgtcagcctcgggcatgggcaggtacacgg-3’.

D78A, fw 5’-tgcccgaggatgccaacaactccctgaacctgggc-3’, rev 5’-gggagttgttggcatcctcgggcatgggcaggtaca-3’.

E76A/D77A, fw 5’-cccatgcccgcggctgacaacaactccctgaacctggg -3’, rev 5’-agttgttgtcagccgcgggcatgggcaggtacacggcca-3’.

E76A/D77A/D78A, fw 5’-atgcccgcggctgccaacaactccctgaacctgggcctg -3’, rev 5’-agttgttggcagccgcgggcatgggcaggtacacggccag-3’.

E90A, fw 5’-gagaagctggcgtacttcttcctcaccgtcttctc -3’, rev 5’-gaagaagtacgccagcttctccaggcccaggttca-3’.

GFP-Ca_V_1.1e-D78A/K162A, -D78A/E90A: To generate these constructs, we employed the same cloning strategy as for D78A but used GFP-Ca_V_1.1e-K162A or -E90A as the template for the PCR reactions.

### CRISPR-mediated genome editing in GLT cells

To generate α_2_δ-1 loss-of-function cell lines, we used the CRISPR/Cas9 system based on the pX458 vector (Addgene plasmid #48138; a gift from Feng Zhang), which co-expresses Cas9, GFP, and a guide RNA. A guide RNA (sgRNA) targeting exon 6 of α_2_δ-1 was designed using the VBC-Score CRISPR design tool (https://www.vbc-score.org/). Cloning of the guide RNA into pX458 was performed following the Zhang lab protocol (https://media.addgene.org/cms/filer_public/95/12/951238bb-870a-42da-b2e8-5e38b37d4fe1/zhang_lab_grna_cloning_protocol.pdf). Briefly, complementary oligonucleotides encoding the sgRNA were annealed and ligated into BbsI-digested pX458. Low-passage GLT cells were plated on 100 mm dishes and transfected with 3.5 μg of the sgRNA-containing pX458 vector using FuGENE HD (Promega). Forty-eight hours after transfection, GFP-positive cells were sorted into 96-well plates containing growth medium using a FACSAria I flow cytometer (BD Biosciences). Untransfected GLTs served as negative controls. Expanded clones were genotyped by PCR using three primers designed to distinguish between wild-type and edited alleles. One amplicon (489 bp) extended beyond the expected indel site, while the second (205 bp) overlapped it. In knockout clones, indel mutations were expected to disrupt the amplification of the shorter, overlapping fragment, while the longer amplicon remained detectable ^24^. The primers used were:

- Forward (Fo): 5′-GCATCTTAACGTGTTTTAAG-3′
- Internal forward (Fi): 5′-CCCACGGACATCTATGAGGGCTG-3′
- Reverse (R): 5′-GAGACAGTTTCTCACTTGC-3′

To characterize insertions and deletions (indels), the 589 bp amplicon spanning exon 6 from selected knockout clones (lacking the 205 bp product) was cloned into the pcDNA3 vector. Multiple pcDNA3 clones were sequenced by Sanger sequencing to identify the specific mutations. Clone C9, which showed a frameshift mutation and superior ability to differentiate into myotubes, was selected for further experiments.

### RT-PCR

Total RNA was isolated from DIV 9–10 GLT and C9 cells using the RNeasy Protect Mini Kit (Qiagen). Reverse transcription was performed with SuperScript II reverse transcriptase (Invitrogen). α_2_δ-1 transcript levels were quantified using a TaqMan assay (Mm00486607_m1, Thermo Fisher Scientific). Absolute transcript copy numbers were calculated using a standard curve generated from PCR products of known concentration, following the procedure described by ^25^. A primer pair was designed to amplify the region spanning exons 6 and 7. The PCR product was purified with the QIAquick Gel Extraction Kit (Qiagen), and DNA concentrations were measured using the Quant-iT PicoGreen dsDNA Assay Kit (Invitrogen). A standard curve was generated from serial dilutions of the purified product (10¹ to 10⁷ molecules) in water containing 1 μg/ml poly-dC-DNA (Midland, TX, USA). Each standard dilution was run in triplicate. Negative controls (no template) were included in each run. Standard curve reliability was assessed across three independent experiments, and average linear regression values were used for quantification.

### Western Blot

Protein lysates were prepared from CRISPR-treated GLT clones at DIV 9–10 as described previously ^26^. Briefly, cells were trypsinized, pelleted by centrifugation, and lysed in ice-cold RIPA buffer using a pestle. Lysates were incubated on ice for 30 minutes, centrifuged, and the supernatant was collected. Protein concentration was determined using a BCA assay (Thermo Scientific). Equal amounts of protein were separated by SDS-PAGE (4–12%) at 196 V and 40 mA for 60 minutes and transferred to PVDF membranes at 25 V and 100 mA for 3 hours at 4°C using a semi-dry transfer system (Roth). Membranes were blocked and incubated overnight at 4°C with mouse anti-α_2_ (1:1,000; clone 20A, MA3-921, Thermo Fisher Scientific) and mouse anti-GAPDH (1:100,000; Santa Cruz Biotechnology). HRP-conjugated secondary antibodies (1:5,000; Pierce) were applied for 1 hour at room temperature. Signals were detected using the ECL SuperSignal West Pico Kit (Thermo Scientific) and visualized with the ImageQuant LAS 4000 system.

### Cell culture and transfection

Dysgenic (α1s-null) myoblasts (GLT cell line) were cultured in growth medium consisting of Dulbecco’s Modified Eagle Medium (DMEM) with 1 g/L glucose, supplemented with 10% fetal calf serum and 10% heat-inactivated horse serum (HS), in culture flasks maintained at 37 °C in a humidified incubator with 10% CO_2_. The cells were then seeded into 35 mm plastic dishes for electrophysiology experiments or onto gelatin and carbon-coated plastic dishes for immunofluorescence analysis. On the second day, and every other day thereafter, the medium was replaced with fusion medium (DMEM supplemented with 2% HS). On the fourth day, the cells were transiently transfected with a specific Ca_V_1.1 construct using FuGeneHD transfection reagent (Promega) following the manufacturer’s protocol. Electrophysiology experiments were conducted on days 7 and 8 after plating, while immunofluorescence staining was carried out on days 9 and 10.

### Immunostaining and Quantification

Methanol-fixed cultures were immunolabeled with the polyclonal rabbit anti-GFP antibody (serum, 1:20000; A6455 Thermofisher Scientific) and the monoclonal mouse anti-DHPR α_2_ antibody (20A, 1:500; MA3-921 Thermofisher Scientific) or with the polyclonal rabbit anti-RyR antibody (1:2000 (AB#5; ^27^) and the monoclonal mouse anti-DHPR α_2_ antibody (20A, 1:500; MA3-921 Thermofisher Scientific) and subsequently fluorescently labeled with Alexa-488 and Alexa-594, respectively, as previously described ^28^. 14-bit images were recorded with a cooled CCD camera (SPOT) and Metaview image processing software. Image composites were arranged in Affinity Designer and Photo (version 2.5.3, Serif (Europe) Ltd.) and non-linear adjustments were performed to correct black level and contrast for the graphical representation. Clusters of GFP-Ca_V_1.1 and α_2_δ-1 were quantified using Image J software (NIH), as previously described ^29^. Briefly, myotubes were selected by a ROI tool and converted to binary images using the intermodes threshold, so that only clusters are included. Using the Analyze Particle function, the numbers of particles larger than 0.2 - 10 μm^2^ in the binary image were counted as clusters. The numbers of clusters per 100 μm^2^ were calculated and are represented in the graphs. To quantify the GFP-Ca_V_1.1/α_2_δ-1 colocalization, the thresholded image was used as a mask on the myotube image to include only the clusters and Pearson ’s coefficients for co-localization were calculated by JaCoP, as previously described ^30^. For each condition, the number of clusters of 30 myotubes from three separate experiments was counted. Graphs and statistical analysis (One-way ANOVA multiple comparison) were performed using GraphPad Prism 10 software package.

### Structure modeling

We predicted the structure of the human WT Ca_V_1.1 α_1_ and α_2_δ subunit complex by making a homology model based on the cryo-EM structure of the rabbit Ca_V_1.1 α_1_ and α_2_δ subunit complex with the VSDs in the up-state and the pore closed (PDB code: 5JGW). Homology modeling for the activated state has been performed using Rosetta and MOE (Molecular Operating Environment, version 2020, Molecular Computing Group Inc, Montreal, Canada). Additionally, we used intermediate and resting state structure model representatives previously characterized in ^12^, as starting points for modelling Ca_V_1.1 α_1_ and α_2_δ-1 complexes and their respective interactions in different IS4 states. The tip of the II-III loop backbone was remodeled using the cryo-EM structure as template, followed by multiple rounds of minimizations and energy optimizations in MOE. The structures were aligned in the membrane using the PPM server and inserted into a plasma membrane consisting of POPC (1-palmitoyl2-oleoyl-sn-glycero-3-phosphocholine) and cholesterol in a 3:1 ratio, using the CHARMM-GUI Membrane Builder ^31^. Water molecules and 0.15 M CaCl_2_ were included in the simulation box. For calcium the standard parameters for calcium-ions were replaced with the multi-site calcium of Zhang et al. This multi-site model has been used to investigate calcium permeation in a number of channels, including type-1 ryanodine receptor ^32–34^.

### Molecular dynamics simulations

All simulations of the complexes were performed with the CHARMM36m force field for the protein, lipids and ions (except for calcium as described above) ^35–37^. The TIP3P water model was used to model solvent molecules ^38^. The system was minimized and equilibrated using the NPT ensemble for a total time of 2 ns with force constraints on the system components being gradually released over six equilibration steps. The temperature was maintained at T = 310 K using the Nosé-Hoover thermostat with the inverse friction constant set to 1.0 ps, and the pressure was maintained semi-isotropically at 1 bar using the Parrinello-Rahman barostat with the period of pressure fluctuations at equilibrium set to 5.0 ps and the compressibility set to 4.5e-5 bar^-1^ ^39,40^. Periodic boundary conditions were used throughout the simulations. Long-range electrostatic interactions were modelled using the particle-mesh Ewald method with a cut-off of 12 Å ^41,42^. The LINCS algorithm was used to constrain bond lengths involving bonds with hydrogen atoms. The simulations were performed using a time step of 2 fs. All simulations were repeated twice for each 200 ns. PyMOL Molecular Graphics System was used to visualize the key interactions and point out differences in the different states (PyMOL Molecular Graphics System, version 2.0, Schrödinger, LLC).

### Dihedral entropies

The dihedral entropies represent an alignment-independent measure of local protein flexibility ^43^. Residue-wise dihedral entropies were calculated by applying the X-Entropy Gaussian kernel density estimator (KDE) to 1D distributions of dihedral angles from the molecular dynamics trajectories. Shannon entropy was then obtained by numerically integrating the KDE-estimated probability density distributions for each dihedral per residue.

### Electrophysiology

Calcium currents were recorded with the patch clamp technique with the cells being voltage-clamped in the whole cell mode. The patch pipettes (borosilicate glass, Harvard Apparatus, Holliston, MA) were prepared to yield a resistance of 2 - 4 MΩ when filled with (mM) 145 Cs-aspartate, 2 MgCl_2_, 10 HEPES, 0.1 Cs-EGTA, and 2 Mg-ATP (pH 7.4 with CsOH). The extracellular bath solution contained (mM) 10 CaCl_2_, 145 tetraethylammonium chloride, and 10 HEPES (pH 7.4 with CsOH).

The recordings were performed with HEKA EPC 10 USB amplifier (Harvard Bioscience Inc.). Data acquisition and command potentials were controlled by PATCHMASTER NEXT (version 1.2, Harvard Bioscience Inc.).

The cells were clamped at a holding potential (V_hold_) of −80 mV, followed by the command potential (V_cmd_) of 500ms, ranging from −60 mV to +80 mV with an increment of 10 mV. Before each main pulse, four leak subtraction pulses were applied with a duration equal to the main pulse, a delay of 10 ms in between the leak pulses, a size of 25% relative to the amplitude of the main pulse and at a holding potential of -120 mV. Between each command potential the cell is repolarized to the holding potential (-80 mV) for a duration of 250 ms. The current-voltage dependence was calculated according to:

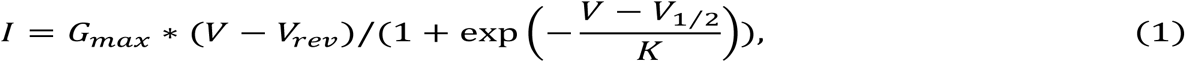

where G_max_ is the maximum conductance of the L-type calcium channels, V_rev_ is the extrapolated reversal potential of the current, V_1/2_ is the potential for half-maximal conductance, and K is the inverse of the slope.

The conductance voltage-dependence was calculated according to the Boltzmann distribution:

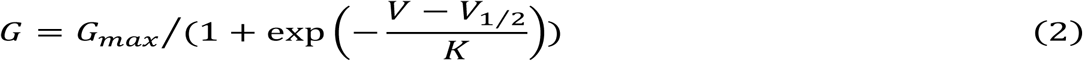

The mean ± SEM for the calcium currents (I_Ca_) sample traces was calculated by selecting the sweep at V_max_ for each recording constituting the dataset. The SEM was calculated for each individual time step.

The time constant of activation (τ_Activation_) was evaluated be fitting the rising phase of the currents at V_max_ with a single exponential function.

### Statistics

Statistical analysis and curve fitting was performed using SigmaPlot (version 12.0.0.182, Grafiti LLC). Figures were prepared in GraphPad Prism (version 10.2.3, GraphPad Software LLC) and Affinity Designer and Photo (version 2.5.3, Serif (Europe) Ltd.). All data are represented as mean ± SEM. Figures presenting current traces the point by point calculated mean ± SEM. When comparing two different data sets, the statistical fit parameters were obtained by using a Student’s t test. In case of non-Gaussian distribution of the data set, tested with a Shapiro-Wilk test, we performed a Mann-Whitney test. If the variances of the data set showed a significant difference, tested with a F-test, a Welch t-test was performed. To compare multiple data sets we performed one-way ANOVA combined with Dunnett’s multiple comparison post hoc test with significance criteria. In case of non-Gaussian distribution of the data set, tested with a Shapiro-Wilk test, we performed a Kruskal-Wallis test with Dunn’s multiple comparison post hoc test. If the variances of the data set showed a significant difference, tested with a Brown-Forsythe test, a Welch ANOVA test with Dunnett’s T3 multiple comparison post hoc test was performed. The significance criteria are as follows *p < 0.05, **p < 0.01, ***p < 0.001 and ****p < 0.0001. The exact p-values for all statistical tests are shown in Tables 1 and 2.

## END NOTES

## Acknowledgements

We thank Matthias Winkler and Pauline Alton for technical help. This research was funded in part by the Austrian Science Fund (FWF) Grant-DOI 10.55776/P35618 to B.E.F. and Grant-DOI 10.55776/P33776 to M.C., and by the Austrian Academy of Sciences APART-MINT postdoctoral fellowship to M.L.F.Q. M.H. was a student of the Ca_V_X PhD program co-funded by FWF (Grant-DOI 10.55776/DOC30) and the Medical University Innsbruck. For open access purposes, the authors have applied a CC BY public copyright license to any author accepted manuscript version arising from this submission.

## Contributions

Experiments were conceived and designed by M.C.H, M.C. and B.E.F. Experiments: Electrophysiology was performed by M.C.H. Structure modeling was performed by M.L.F.Q. Generation of expression plasmids and of the GLT-C9 cell line were planned and supervised by M.C. and performed by H.E.A. Immunofluorescence data were analyzed and figures prepared by N.C, B.E.F., M.C.H., and M.C. Coworkers were trained and the project was supervised by M.C. and B.E.F. The paper was written by B.E.F. with input from all authors.

